# Extracellular NAD(P) activates systemic acquired resistance through LecRK-VI.2-mediated phosphorylation of NPR1

**DOI:** 10.64898/2026.06.02.729676

**Authors:** Cheng Liu, Qingcai Liu, Shweta Chhajed, Mingxi Zhou, Fiona M. Harris, Xudong Zhang, Sixue Chen, Zhonglin Mou

## Abstract

Systemic acquired resistance (SAR) is a long-lasting, broad-spectrum immune response induced in distal tissues by signals generated at primary infection sites. Although numerous mobile immune signals have been implicated in SAR, how these signals are perceived and mechanistically coupled to transcriptional reprogramming in systemic tissues remains poorly understood. Extracellular NAD(P) [eNAD(P)] functions as a key integrative SAR signal that activates immunity through the plasma membrane-localized lectin receptor kinase LecRK-VI.2 and the master immune coactivator NONEXPRESSOR OF PATHOGENESIS-RELATED GENES1 (NPR1). However, the mechanism linking eNAD(P) perception to activation of NPR1 has remained unknown. Here, we show that LecRK-VI.2 constitutively associates with NPR1 and directly phosphorylates NPR1 at T359 and likely S356 upon eNAD(P) perception. NADP^+^-induced phosphorylation of NPR1 occurs rapidly in vivo and requires LecRK-VI.2. Nonphosphorylatable NPR1 variants abolish eNADP^+^-induced local and systemic immunity as well as biologically induced SAR, whereas phosphomimetic variants retain NPR1 function. Mechanistically, LecRK-VI.2-mediated phosphorylation promotes NPR1 interaction with TGACG-binding transcription factors (TGAs) and the Mediator subunit MED15, thereby enhancing assembly of a transcriptional activation complex required for defense gene expression. We further demonstrate that NPR1 facilitates TGA-MED15 association in a phosphorylation-dependent manner. Together, these findings establish a receptor-to-coactivator signaling mechanism that directly links extracellular immune signal perception to transcriptional activation. This work closes a major mechanistic gap in the SAR signaling pathway and reveals receptor-mediated coactivator activation as a mechanism for rapid conversion of extracellular immune cues into coordinated transcriptional outputs during systemic immunity.

## Introduction

Multicellular eukaryotes have evolved sophisticated immune systems to counter microbial infections. While animals deploy both innate and adaptive immunity, plants rely exclusively on germline-encoded receptors, including cell-surface pattern recognition receptors (PRRs) and intracellular nucleotide-binding domain leucine-rich repeat receptors (NLRs), which mediate pattern- and effector-triggered immunity (PTI and ETI), respectively (Jones and Dangl, 2006; Jones et al., 2024; Zipfel, 2014). Activation of PTI and ETI at infection sites leads to the production of mobile immune signals that travel to distal tissues to establish systemic acquired resistance (SAR), a long-lasting and broad-spectrum defense response (Fu and Dong, 2013; Kachroo and Kachroo, 2020).

A diverse set of candidate SAR mobile signals has been identified, including DEFECTIVE IN INDUCED RESISTANCE1 (Maldonado et al., 2002), salicylic acid (SA) and its derivative methyl SA (Park et al., 2007; Shulaev et al., 1997; Wang et al., 2018), dehydroabietinal (Chaturvedi et al., 2008), azelaic acid (AzA) (Jung et al., 2009), glycerol-3-phosphate (G3P) (Chanda et al., 2011), pipecolic acid (Pip) and its derivative N-hydroxy-Pip (NHP) (Chen et al., 2018; Hartmann et al., 2018; Navarova et al., 2012), extracellular NAD(P) [eNAD(P)] (Li et al., 2023), monoterpenes (Riedlmeier et al., 2017), and phased small RNAs (Shine et al., 2022). In addition, nitric oxide and reactive oxygen species (ROS) have been shown to play significant roles in SAR (Alvarez et al., 1998; Wang et al., 2014). Emerging evidence suggests that SAR mobile signals initiate a signal amplification loop that enhances ROS production in systemic tissues (Alvarez *et al*., 1998; Riedlmeier *et al*., 2017; Wang *et al*., 2014; Wang *et al*., 2018; Wenig et al., 2019; Zottini et al., 2007), which in turn triggers the accumulation of eNAD(P) (Li *et al*., 2023). Preventing systemic accumulation of eNAD(P) compromises SAR (Li *et al*., 2023), whereas exogenous NAD(P) can restore systemic immunity in mutants defective in multiple SAR signaling pathways (Wang et al., 2019). Although these findings establish eNAD(P) as a convergence point that integrates various SAR signals in systemic tissues (Li *et al*., 2023; Liu et al., 2024), how eNAD(P) perception is coupled to nuclear gene activation remains unclear.

Two plasma membrane-localized legume-like (L-type) lectin receptor kinases (LecRKs), LecRK-I.8 and LecRK-VI.2, have been identified as eNAD(P) receptors (Wang *et al*., 2019; Wang et al., 2017). Loss of these receptors impairs eNAD(P)-induced immunity and SAR (Li et al., 2024; Wang *et al*., 2019; Wang *et al*., 2017), underscoring their importance in systemic defense. However, the mechanisms by which these cell-surface receptors transduce eNAD(P) perception into intracellular signaling and transcriptional reprogramming remains undefined. As receptor kinases with active catalytic domains (Bouwmeester and Govers, 2009; Singh et al., 2013; Wang *et al*., 2017), LecRKs are well positioned to directly relay eNAD(P) signals through phosphorylation of cytosolic targets.

NONEXPRESSOR OF PATHOGENESIS-RELATED GENES1 (NPR1) is a master transcriptional coactivator that regulates immune gene expression through interaction with TGACG-binding factors (TGAs) and other transcriptional regulators (Després et al., 2000; Zavaliev and Dong, 2024; Zhang et al., 1999; Zhou et al., 2000). NPR1 resides predominantly in the cytosol prior to activation and accumulates in the nucleus upon immune induction (Kinkema et al., 2000). Because NPR1 acts downstream of eNAD(P) in the SAR signaling pathway (Wang *et al*., 2019; Zhang and Mou, 2009), we hypothesized that eNAD(P) perception may directly engage NPR1 to couple extracellular signal perception to transcriptional activation.

Here, we show that LecRK-VI.2 physically associates with NPR1 in an uninduced state and directly phosphorylates NPR1 at T359 and likely S356 upon eNAD(P) perception. This modification is required for NPR1 interactions with TGAs and the Mediator subunit MED15, enabling assembly of a transcriptional activation complex and induction of defense gene expression. These findings establish a receptor-to-coactivator signaling mechanism that directly link extracellular signal perception to transcriptional activation, providing a mechanistic framework for how systemic immune signals are converted into coordinated transcriptional outputs in SAR.

## Results

### LecRK-VI.2 interacts with NPR1

To investigate whether LecRK-VI.2 interacts with NPR1, we first performed pairwise yeast two-hybrid (Y2H) assays using the intracellular kinase domains (KDs) of LecRK-VI.2 (LecRK-VI.2KD) and LecRK-I.8 (LecRK-I.8KD) as bait. While yeast cells expressing LecRK-VI.2KD or LecRK-I.8KD with NPR1 as prey grew on synthetic dextrose (SD) double dropout (DDO) (-Trp-Leu) medium, only those expressing LecRK-VI.2KD as bait and NPR1 as prey grew on SD quadruple dropout (QDO) (-Trp-Leu-His-Ade) medium (Figure 1A). These results indicate that LecRK-VI.2KD, but not LecRK-I.8KD, physically interacts with NPR1 in yeast.

**Figure 1.**
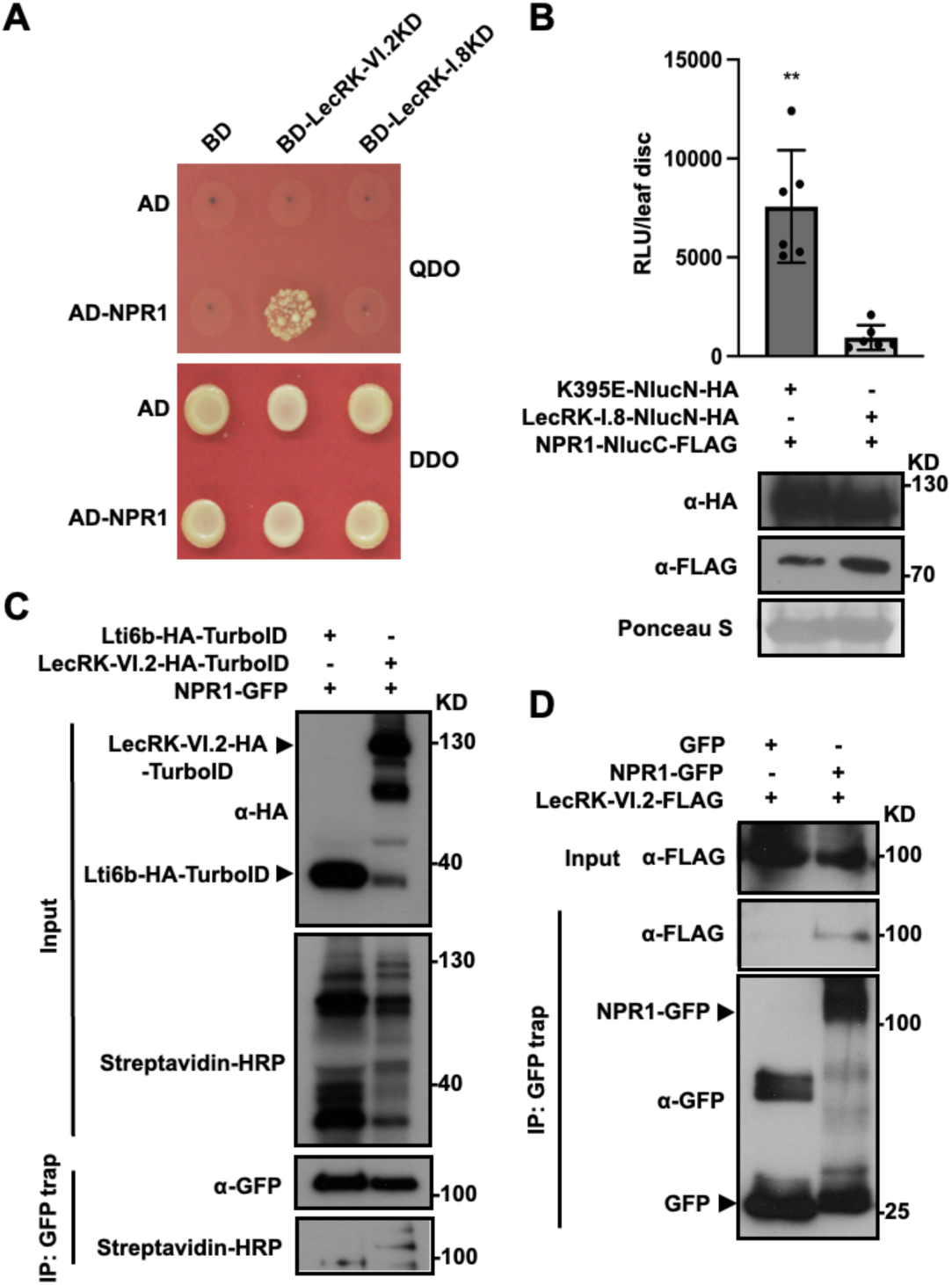
LecRK-VI.2 forms a complex with NPR1. **(A)** Interaction between LecRK-VI.2KD and NPR1 in yeast. BD, DNA binding domain; AD, activation domain. **(B)** Split Nano luciferase complementation assay of the interaction between LecRK-VI.2 and NPR1 in *N. benthamiana*. Ponceau S staining of Rubisco was used as the loading control. RLU, relative light unit. Bars represent means ± standard deviation (SD) (n = 6). Asterisks denote a significant difference (**p < 0.01; Student’s t-test). **(C)** TurboID-based proximity labeling assessing the association between LecRK-VI.2 and NPR1 in transgenic *Arabidopsis*. Total protein was extracted from biotin-treated leaves and subjected to immunoprecipitation with GFP-trap magnetic beads, followed by immunoblotting with streptavidin-HRP and anti-GFP antibody. **(D)** Co-immunoprecipitation assay of the interaction between LecRK-VI.2 and NPR1 in *N. benthamiana*. Immunoprecipitation was conducted using GFP-trap magnetic beads, followed by immunoblotting with anti-FLAG and anti-GFP antibodies. Molecular mass markers in kilo Daltons (KD) are indicated on the right in (**B** to **D**). The experiments were repeated at least three times with similar results.

To confirm this interaction, we conducted split Nano luciferase (Nluc) complementation assays (Wang et al., 2020). Since LecRK-VI.2 induces cell death when transiently expressed in *Nicotiana benthamiana* (Li *et al*., 2024), we used its kinase-dead variant, lecrk-VI.2^K395E^ (K395E). Strong luciferase activity was detected when K395E and NPR1 were co-expressed as NlucN and NlucC translational fusions, respectively (Figure 1B). By contrast, replacing K395E with LecRK-I.8 led to a significant reduction in luciferase activity, despite verified expression of LecRK-I.8-NlucN-HA and NPR1-NlucC-FLAG in the negative control (Figure 1B). This result supports physical association between LecRK-VI.2 and NPR1.

To determine whether LecRK-VI.2 associates with NPR1 *in vivo*, we employed TurboID-based proximity labeling (Branon et al., 2018). Stable transgenic *Arabidopsis* plants co-expressing NPR1-GFP with either LecRK-VI.2-HA-TurboID or Lti6b-HA-TurboID (negative control) were generated. Significant biotinylation of NPR1-GFP was observed only in plants co-expressing LecRK-VI.2-HA-TurboID (Figure 1C), supporting the *in vivo* association between LecRK-VI.2 and NPR1. Furthermore, co-immunoprecipitation (co-IP) assays showed that LecRK-VI.2-FLAG was co-IPed with NPR1-GFP, but not GFP alone, in *N. benthamiana* (Figure 1D), confirming that LecRK-VI.2 physically interacts with NPR1.

### LecRK-VI.2 phosphorylates NPR1 *in vitro* and *in vivo*

Since receptor-like kinases often regulate signaling by phosphorylating their interactors, we tested whether LecRK-VI.2 phosphorylates NPR1 using *in vitro* kinase assays. Purified MBP-tagged LecRK-VI.2KD (MBP-LecRK-VI.2KD) efficiently phosphorylated superfolder GFP (sfGFP)- and His-tagged NPR1 (sfGFP-NPR1-His), but not sfGFP-His, as detected by an anti-phosphothreonine (pThr) antibody (Figure 2A). To determine whether NPR1 phosphorylation by LecRK-VI.2 is specific, we included BAK1KD, the kinase domain of BAK1, a leucine-rich repeat receptor kinase known to interact with LecRK-VI.2 (Wang *et al*., 2019). Although GST-BAK1KD interacted with NPR1 in yeast, it failed to phosphorylate sfGFP-NPR1-His *in vitro* (Supplemental Figures 1A and 1B), demonstrating that LecRK-VI.2KD is specifically responsible for NPR1 phosphorylation. To identify the phosphorylation sites, we subjected LecRK-VI.2KD-treated and BAK1KD-treated NPR1 proteins to liquid chromatography-tandem mass spectrometry (LC-MS/MS). This analysis identified S356, T359, and T373 as confident *in vitro* phosphorylation sites on NPR1 by LecRK-VI.2KD, with S571 as a candidate site (Supplemental Figures 2A-2D). Consistent with the kinase assay results, no phosphorylation sites were detected on NPR1 following BAK1KD treatment.

**Figure 2.**
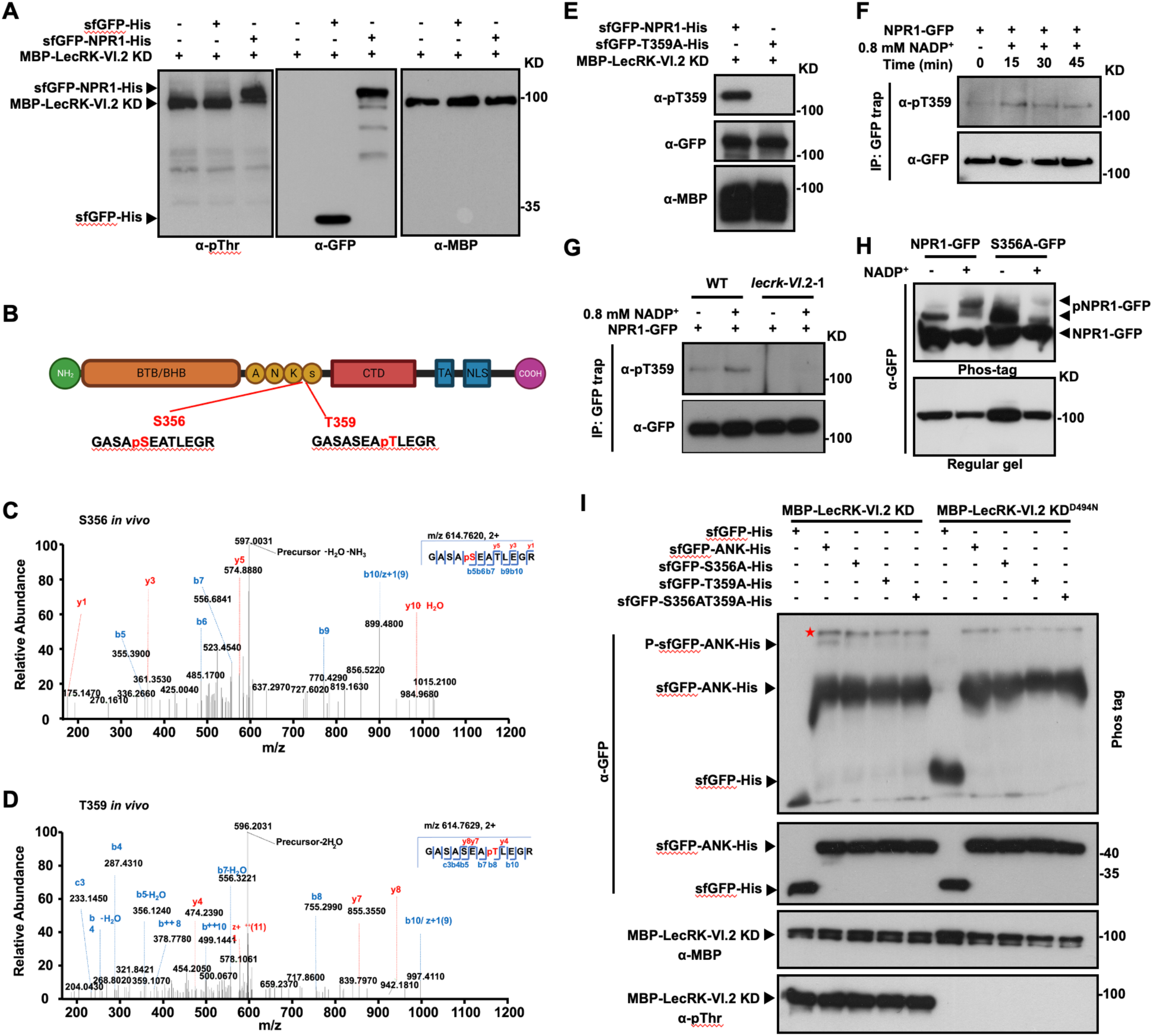
NPR1 is phosphorylated by LecRK-VI.2 *in vitro* and *in vivo*. **(A)** Phosphorylation of NPR1 by LecRK-VI.2KD *in vitro*. Phosphorylation was detected using anti-pThr antibody. The abundance of the recombinant proteins was determined by immunoblotting with anti-GFP and anti-MBP antibodies. **(B)** Schematic diagram of the NPR1 protein motifs with the confident *in vivo* phosphorylation sites identified by LC-MS/MS. **(C** and **D)** LC-MS/MS spectra of peptides harboring pS356 (**C**) and pT359 (**D**) identified in NADP^+^-treated transgenic plants expressing LecRK-VI.2-HA-TurboID and NPR1-GFP. **(E)** Specificity of the anti-pT359 antibody. The abundance of the recombinant proteins was determined by immunoblotting with anti-GFP and anti-MBP antibodies. Phosphorylation at T359 was analyzed using the anti-pT359 antibody. **(F)** NADP^+^-induced T359 phosphorylation. Leaves were infiltrated with 0.8 mM NADP^+^ and collected at the indicated times. Immunoprecipitation was performed using GFP-trap magnetic beads, followed by immunoblotting, first with the anti-pT359 antibody, then with anti-GFP antibody. **(G)** Dependence of NADP^+^-induced T359 phosphorylation on LecRK-VI.2. Leaves were infiltrated with 0.8 mM NADP^+^ and collected at 30 min. Immunoprecipitation was performed using GFP-trap magnetic beads, followed by immunoblotting with anti-pT359 and anti-GFP antibodies. WT, wild type. **(H)** Phos-tag gel analysis of NADP^+^-induced phosphorylation of S356 *in vivo*. Leaves were infiltrated with 0.8 mM NADP^+^ and collected at 30 min. Total proteins were extracted, separated in a phos-tag gel or a regular SDS-PAGE gel, and probed with anti-GFP antibody. **(I)** Phos-tag gel analysis of S356 and T359 phosphorylation of the NPR1 ankyrin repeat (ANK) domain by LecRK-VI.2 *in vitro*. Recombinant proteins of the NPR1 ANK domain (AA 230-360) and the corresponding nonphosphorylatable variants were purified and incubated with MBP-LecRK-VI.2KD or MBP-LecRK-VI.2KD^D494N^. Proteins were separated in a phos-tag or a regular SDS-PAGE gel, and probed with anti-GFP, anti-MBP, and anti-pThr antibodies. The red star indicates potentially phosphorylated sfGFP-ANK-His by unknown kinases in *E. coli*. Molecular mass markers in kilo Daltons (KD) are indicated on the right in (**A** and **E** to **I**). The experiments in (**A** and **E** to **I**) were repeated at least twice with similar results.

To examine whether eNAD(P) induces NPR1 phosphorylation at these sites, we performed LC-MS/MS on NPR1-GFP immunoprecipitated from NADP^+^-treated *Arabidopsis* plants. This analysis identified S356 and T359 as confident *in vivo* phosphorylation sites on NPR1, with T572, S573, and S574 as candidate sites (Figures 2B-2D; Supplementa Figure 2A). To confirm that LecRK-VI.2 is responsible for phosphorylation at these sites, we generated phospho-specific antibodies against key residues. The anti-pT359 antibody specifically recognized MBP-LecRK-VI.2KD-treated sfGFP-NPR1-His, but not the sfGFP-T359A-His mutant (Figure 2E), verifying its specificity for phospho-T359.

Using this antibody, we examined NADP^+^-induced phosphorylation of NPR1 *in vivo*. Immunoprecipitated NPR1-GFP from NADP^+^-treated *NPR1-GFP/npr1-3* plants was analyzed by immunoblotting with anti-pT359. NADP^+^ induced T359 phosphorylation as early as 15 min post-treatment (Figure 2F). Importantly, T359 phosphorylation was abolished in *NPR1-GFP/lecrk-VI.2-1* plants (Figure 2G), demonstrating that LecRK-VI.2 is required for NADP^+^-induced phosphorylation of NPR1 *in vivo*. To determine whether S356 is also phosphorylated in response to eNAD(P), we analyzed NPR1 phosphorylation using Phos-tag gels in *S356A-GFP/npr1-3* transgenic plants. While NADP^+^ treatment induced an upshifted band corresponding to phosphorylated NPR1-GFP, this shift was dramatically reduced in the S356A-GFP mutant (Figure 2H), indicating that S356 is required for NADP^+^-induced NPR1 phosphorylation.

To further assess the contribution of S356 and T359, we analyzed LecRK-VI.2-mediated phosphorylation of the sfGFP- and His-tagged NPR1 ankyrin repeat domain (AA 230-360; sfGFP-ANK-His) and its nonphosphorylatable variants (S356A, T359A, and S356A/T359A) using Phos-tag gels. Strikingly, each single mutation nearly abolished the phosphorylation-dependent mobility shift, and the double mutant showed a similar effect (Figure 2I), indicating that both S356 and T359 are individually required for efficient LecRK-VI.2-mediated phosphorylation of NPR1. Together, these results provide orthogonal biochemical and genetic evidence that eNAD(P) induces LecRK-VI.2-mediated phosphorylation of NPR1 at T359 and likely S356 *in vivo*, establishing NPR1 as a key downstream target of LecRK-VI.2 in eNAD(P) signaling.

### S356/T359 phosphorylation is required for NPR1 function

To determine the biological function of eNAD(P)-induced, LecRK-VI.2-mediated phosphorylation of NPR1, we compared NADP^+^-induced local and systemic immunity, as well as biologically induced SAR, in *NPR1-GFP*/*npr1-3*, *S356A/T359A-GFP*/*npr1-3*, and *S356D/T359D*/*npr1-3* plants. While NPR1-GFP and S356D/T359D-GFP fully complemented the *npr1-3* mutant, S356A/T359A-GFP failed to do so (Figures 3A-3C). Consistently, NADP^+^-induced *PR1* gene expression and PR1 protein accumulation were significantly reduced in *S356A/T359A-GFP*/*npr1-3* plants but remained unaffected in *S356D/T359D-GFP*/*npr1-3* transgenic plants (Supplemental Figures 3A and 3B; Figures 3D and 3E). Furthermore, our LC-MS/MS analysis, along with prior research (Yao et al., 2023), identified T572, S573, and S574 as potential *in vivo* phosphorylation sites (Supplemental Figure 2A). However, substitution of S/T571-576 with six alanine residues (6A) did not affect NPR1 function in NADP^+^-induced immunity and SAR (Supplemental Figures 4A-4D). Together, these results demonstrate that phosphorylation of S356/T359 is essential for NPR1 function in eNAD(P) signaling and SAR.

**Figure 3.**
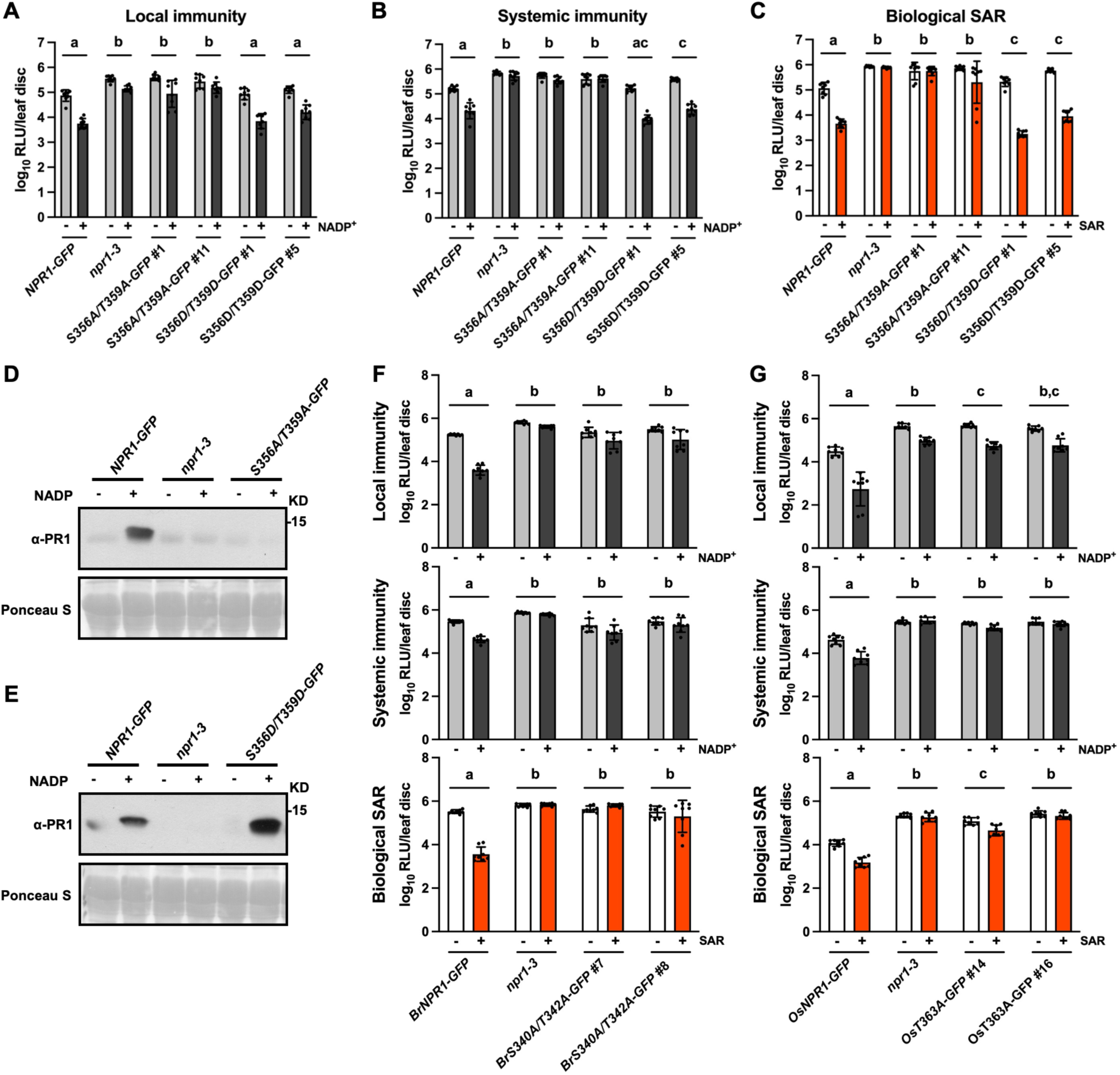
NPR1 function depends on phosphorylation at S356/T359. **(A)** NADP^+^-induced local immunity in the indicated genotypes. Two leaves on each plant were infiltrated with 0.8 mM NADP^+^ or water. Four hr later, the infiltrated leaves were inoculated with a *Psm-lux* suspension (OD_600_ = 0.001). **(B)** NADP^+^-induced systemic immunity in the indicated genotypes. Three leaves on each plant were infiltrated with 0.8 mM NADP^+^ or water. Four hr later, two systemic leaves were inoculated with a *Psm*-lux suspension (OD_600_ = 0.001). **(C)** Biological induction of SAR in the indicated genotypes. Three lower leaves on each plant were infiltrated with *Psm* (OD_600_= 0.002) or 5 mM MgCl_2_. Two days later, two systemic leaves were inoculated with a *Psm*-lux suspension (OD_600_ = 0.001). In (**A** to **C**), samples were collected 2.5 days later. Bars represent means ± SD (n = 8). Different letters denote significant differences (p < 0.05; two-way ANOVA). **(D** and **E)** NADP^+^-induced PR1 protein accumulation in *npr1-3* and *npr1-3* complementation lines expressing NPR1-GFP or its phospho-mutants. Leaves were treated with 0.8 mM NADP^+^ for 24 hr. Total proteins were extracted and analyzed by immunoblotting with anti-PR1 antibody. Ponceau S staining of Rubisco was used as the loading control. **(F** and **G)** NADP^+^-induced local and systemic immunities as well as biological induction of SAR in *npr1-3* and *npr1-3* complementation lines expressing *BrNPR1*, *OsNPR1*, or their phospho-mutants. The experiments were conducted as in (**A**), (**B**), and (**C**). Bars represent means ± SD (n = 8). Different letters denote significant differences (p < 0.05; two-way ANOVA). The experiments were repeated at least three times with similar results.

Importantly, residues corresponding to S356 and T359 are highly conserved among NPR1 orthologs in various plant species (Supplemental Figures5A and 5B) (Jia et al., 2023), suggesting a conserved functional role. To investigate this, we assessed the function of nonphosphorylatable NPR1 ortholog variants from *Brassica rapa* (BrNPR1) and *Oryza sativa* (OsNPR1), both of which have been shown to complement the *Arabidopsis npr1* mutant (Potlakayala et al., 2007; Yuan et al., 2007). Specifically, BrNPR1 S340 and OsNPR1 T363, corresponding to *Arabidopsis* NPR1 S356 and T359, respectively (Supplemental Figure 5C), were substituted with alanine. Additionally, BrNPR1 T342, adjacent to the conserved T359-equivalent residue (Supplemental Figure 5C), was replaced with alanine. Compared to wild-type BrNPR1 and OsNPR1, the nonphosphorylatable mutations, BrS340A/T342A and OsT363A, reduced yeast growth in Y2H assays with TGA transcription factors on QDO medium (Supplemental Figures 5D and 5E). BrS340A/T342A impaired interaction with TGA1, TGA2, and TGA3 (Supplemental Figure 5D), whereas OsT363A specifically affected interaction with TGA1 (Supplemental Figure 5E). Consistent with this, these mutants failed to complement the *npr1-3* defects in NADP^+^-induced resistance and biological SAR (Figures 3F and 3G). Together, these results demonstrate that phosphorylation of these conserved serine and/or threonine residues is important for NPR1 function across species.

### S356/T359 phosphorylation is required for NPR1-TGA interactions

To investigate the mechanisms underlying eNAD(P) signaling, we first tested whether eNAD(P) induces NPR1 dissociation from LecRK-VI.2. We treated *N. benthamiana* leaves transiently co-expressing LecRK-VI.2-FLAG and NPR1-GFP with NADP^+^ and analyzed their association via co-IP. While similar amounts of NPR1-GFP were pulled down with GFP-trap beads from treated and untreated samples, less LecRK-VI.2-FLAG was co-IPed from NADP^+^-treated samples (Supplemental Figure 6A), indicating that NADP^+^ induces NPR1 dissociation from LecRK-VI.2. To further explore this, we performed a time-course experiment using *Arabidopsis NPR1-GFP*/*LecRK-VI.2-HA-TurboID* transgenic plants. NADP^+^ induced biphasic changes in NPR1-GFP and LecRK-VI.2-HA-TurboID association, with a sharp decrease at 30 min post-treatment, followed by a gradual increase surpassing basal levels by 480 min (Supplemental Figure 6B).

Given that LecRK-VI.2 phosphorylates NPR1 within 15 min of NADP^+^ treatment (Figure 2F), and that dissociation from the receptor complex occurs thereafter (Supplemental Figure 6B), we hypothesized that NPR1 phosphorylation may facilitate its interaction with TGA transcription factors. To test this, we analyzed NADP^+^-induced NPR1-TGA interactions in *Arabidopsis NPR1-GFP*/*TGA-HA-TurboID* transgenic plants. Co-IP assays showed a significant increase in NPR1-GFP binding to TGA1 and TGA3 at 60 min post-treatment (Supplemental Figures 6C and 6D), occurring sequentially after NPR1 phosphorylation (15 min) and dissociation from LecRK-VI.2 (30 min) (Figure 2F; Supplemental Figure 6B). The tight correlation among these events suggested that LecRK-VI.2-mediated phosphorylation of NPR1 might be a prerequisite for its interaction with TGAs.

To test this, we first examined the effect of nonphosphorylatable mutations on NPR1-TGA3 interaction using Y2H assays. Compared to wild-type NPR1, the T359A and S356A/T359A mutants significantly reduced and completely suppressed, respectively, yeast growth on QDO medium (Supplemental Figure 7), suggesting that phosphorylation at these sites additively promotes NPR1-TGA binding. Consistently, yeast cells expressing NPR1 or the phosphomimetic S356D/T359D mutant as bait, together with TGA2, TGA3, TGA5, or TGA6 as prey, grew normally on QDO medium (Figure 4A). In contrast, the S356A/T359A mutant failed to support yeast growth (Figure 4A). Interestingly, while wild-type NPR1 did not interact with TGA1 in yeast, S356D/T359D did (Figure 4A), suggesting that S356/T359 phosphorylation may enable NPR1-TGA1 interaction in yeast.

**Figure 4.**
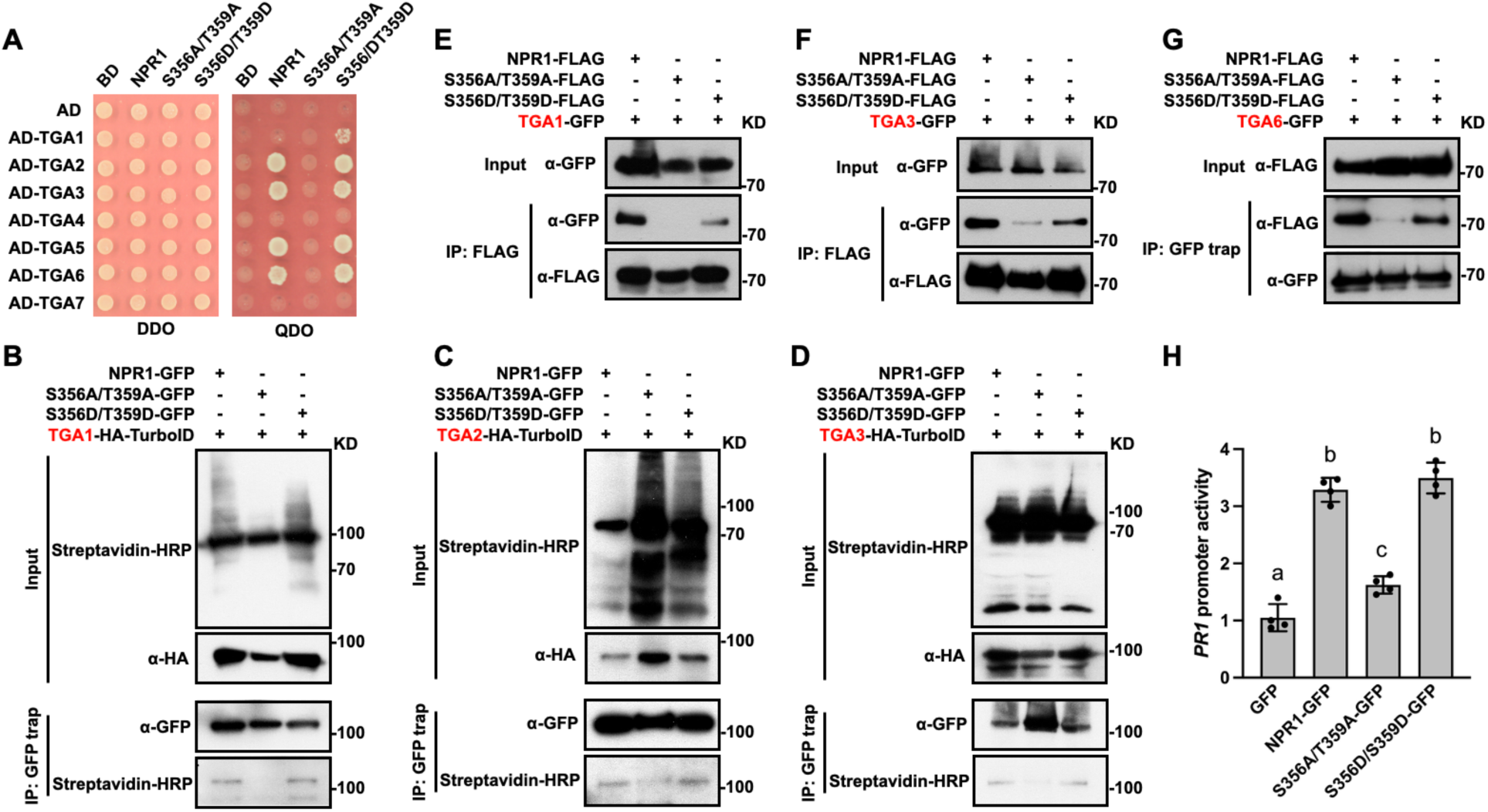
Phosphorylation of NPR1 at S356/T359 enables interaction with TGAs. **(A)** Interactions between NPR1 or its phospho-mutants and TGAs in yeast. The interactions were determined by growth on QDO medium. **(B** to **D)** TurboID-based proximity labeling assays of the associations between NPR1 or its phospho-mutants and TGA1 (**B**), TGA2 (**C**), or TGA3 (**D**) in transgenic *Arabidopsis*. Immunoprecipitation was performed using GFP-trap magnetic beads, followed by immunoblotting with streptavidin-HRP and anti-GFP antibody. **(E** to **G)** Co-immunoprecipitation assays of the interactions between NPR1 or its phospho-mutants and TGA1 (**E**), TGA3 (**F**) or TGA6 (**G**) in *N. benthamiana*. Immunoprecipitation was conducted using anti-FLAG antibody-coupled agarose beads (**E** and **F**) or GFP-trap magnetic beads (**G**), followed by immunoblotting with anti-GFP and anti-FLAG antibodies. **(H)** Induction of the *PR1* promoter by GFP, NPR1-GFP, or its phospho-mutants. Agrobacteria carrying the *pPR1:DUAL-LUC* reporter and the indicated effector construct were co-infiltrated into *N. benthamiana* leaves. Samples were collected two days later for FLUC and RLUC activity assays. *PR1* promoter activity was indicated by the ratio of F-LUC and R-LUC activities for each effector, normalized to free GFP, which was set to 1. Bars represent means ± SD (n = 4). Different letters denote significant differences (p < 0.01; one-way ANOVA). The molecular mass markers in kilo Daltons (KD) are indicated on the right in (**B**-**G**). The experiments were repeated at least twice with similar results.

To examine these interactions *in planta*, we applied TurboID-based proximity labeling in *Arabidopsis* transgenic lines co-expressing NPR1-GFP or its phospho-mutants with TGA1-HA-TurboID, TGA2-HA-TurboID, or TGA3-HA-TurboID. The S356A/T359A mutations, but not S356D/T359D, significantly reduced NPR1-GFP biotinylation (Figures 4B-4D), supporting the requirement of S356/T359 phosphorylation for NPR1-TGA proximity *in vivo*.

We next performed co-IP assays using *N. benthamiana*. Consistent with the Y2H and proximity labeling results, less TGA1-GFP and TGA3-GFP were co-IPed with S356A/T359A-FLAG compared to NPR1-FLAG or S356D/T359D-FLAG (Figures 4E and 4F), and similarly, less S356A/T359A-FLAG was co-IPed with TGA6-GFP (Figure 4G), confirming that the nonphosphorylatable mutations impair NPR1-TGA interactions. Although the S356D/T359D mutant retained association with TGAs, the interaction was reduced relative to wild-type NPR1 (Figures 4E-4G), consistent with the imperfect phosphomimetic nature of aspartic acid (Dephoure et al., 2013).

Since NPR1 functions as a transcriptional coactivator for TGAs (Rochon et al., 2006), we hypothesized that disrupting NPR1-TGA interaction through nonphosphorylatable mutations might compromise its coactivator activity. To test this, we used a *pPR1:DUAL-LUC* reporter assay in *N. benthamiana* (Kumar et al., 2022). S356A/T359A-GFP failed to fully activate the *PR1* promoter, a direct target of NPR1, whereas S356D/T359D-GFP exhibited wild-type levels of transcriptional activity (Figure 4H). These results indicate that phosphorylation at S356/T359 is required for NPR1 coactivator function.

### NPR1 and TGAs interact with MED15

The Mediator complex is a large, multiprotein transcriptional coactivator in eukaryotes that transduces signals from pathway-specific transcription factors to RNA polymerase II (Richter et al., 2022). Subunits MED14 and MED16 are required for eNAD(P) signaling (Zhang et al., 2012; Zhang et al., 2013). MED15 is also essential, as NADP^+^-induced local and systemic immunity as well as PR1 protein accumulation were impaired in the *nrb4-3* mutant (a *med15* allele) (Canet et al., 2012), similar to the *tag2/3/5/6* and *npr1-3* mutants (Figures 5A-5C) (Cao et al., 1997; Kesarwani et al., 2007). Furthermore, MED14, Med15, and MED16 have been shown to be in proximity to NPR1 (Powers et al., 2024), suggesting that NPR1 and/or TGAs might recruit the Mediator complex through these subunits. To test this hypothesis, we first examined whether NPR1 interacts directly with MED14, MED15, or MED16 using Y2H assays. Yeast cells expressing MED15 as bait and NPR1 as prey grew on TDO medium supplemented with 5 mM 3-amino-1,2,4-triazole (3-AT), whereas those expressing MED14 or MED16 as bait did not grow (Figure 5D). These results indicate that NPR1 interacts specifically with MED15 in yeast.

**Figure 5.**
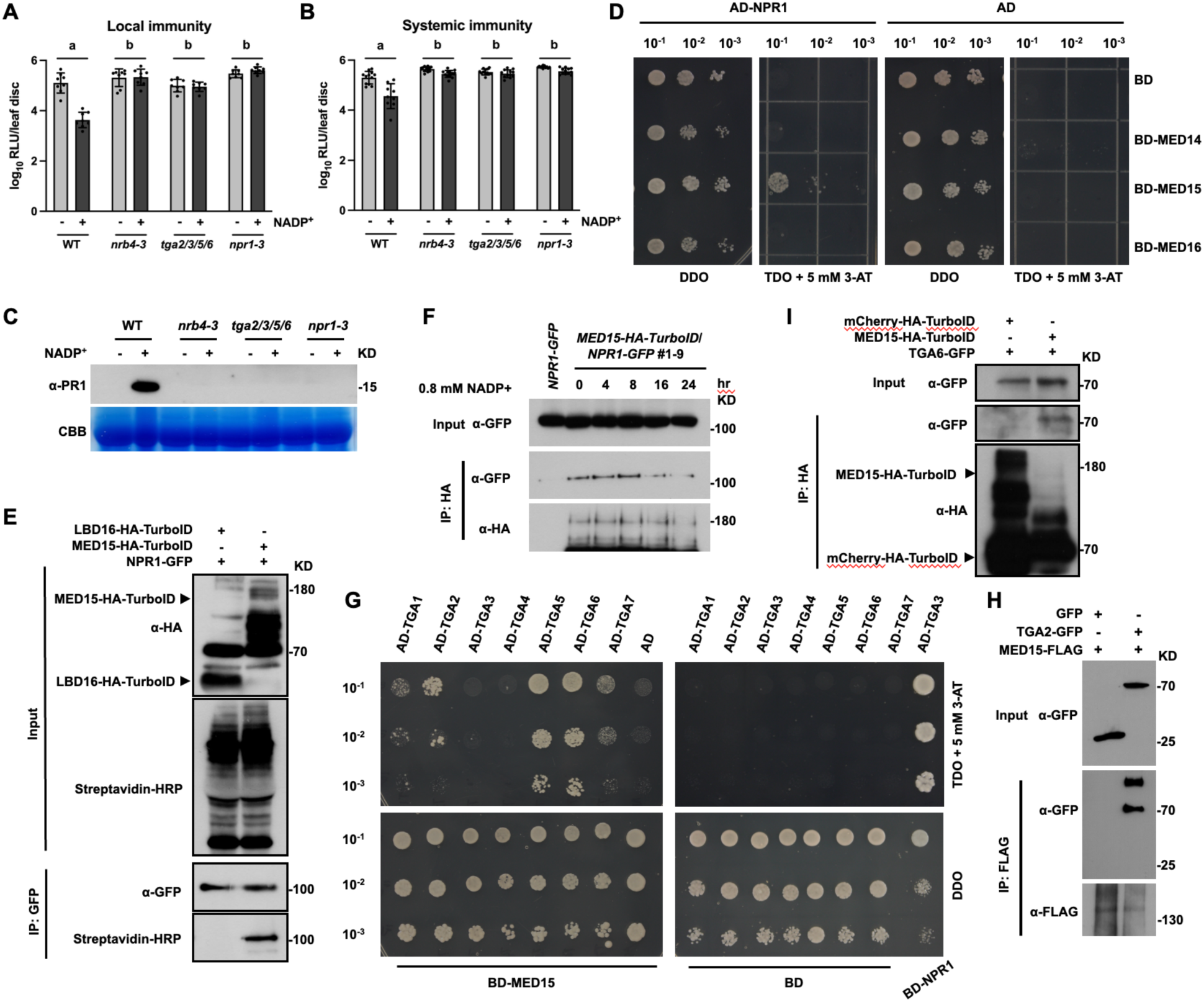
NPR1 and TGAs associate with MED15. **(A** and **B)** NADP^+^-induced local (**A**) and systemic (**B**) immunity in the indicated genotypes. Bars represent means ± SD (n = 8-12). Different letters denote significant differences (p < 0.05; two-way ANOVA). **(C)** NADP^+^-induced PR1 protein accumulation in the indicated genotypes. Coomassie brilliant blue (CBB) staining of Rubisco was used as the loading control. **(D)** Yeast two-hybrid assay examining the interaction between NPR1 and Mediator subunits. The full-length *MED14*, *MED15*, and *MED16* were cloned into the bait vector pGBKT7, and the full-length *NPR1* was cloned into the prey vector pGADT7. The bait and prey vectors were co-transformed into the yeast strain AH109, and the resulting yeast cells were grown on DDO medium. Interaction was determined by growth on triple dropout (TDO) (-Trp-Leu-His) medium supplemented with 5 mM 3-amino-1,2,4-triazole (3-AT). **(E)** TurboID-based proximity labeling examining the association between NPR1 and MED15 in transgenic *Arabidopsis* expressing NPR1-GFP and MED15-HA-TurboID or LBD16-HA-TurboID. Total protein was extracted from biotin-treated leaves and subjected to immunoprecipitation with GFP-trap magnetic beads, followed by immunoblotting with streptavidin-HRP and anti-GFP antibody. **(F)** Co-immunoprecipitation assays examining the interaction between NPR1 and MED15 in transgenic *Arabidopsis*. Leaves of four-week-old double transgenic plants expressing NPR1-GFP and MED15-HA-TurboID were treated with 0.8 mM NADP^+^ for the indicated times. Immunoprecipitation was conducted using anti-HA magnetic beads. The precipitated proteins were analyzed by immunoblotting with anti-GFP and anti-HA antibodies. **(G)** Yeast two-hybrid assays showing the interactions between TGAs and MED15. The full-length *MED15* was cloned into the bait vector pGBKT7, and the full-length *TGAs* were in the prey vector pGADT7. The bait and prey vectors were co-transformed into the yeast strain AH109, and the resulting yeast cells were grown on DDO medium. Interaction was determined by growth on TDO medium supplemented with 5 mM 3-AT. **(H)** Co-immunoprecipitation assays of the interactions between MED15 and TGA2 in *N. benthamiana*. MED15-FLAG was transiently co-expressed with TGA2-GFP or GFP in *N. benthamiana*. Immunoprecipitation was carried out using anti-FLAG antibody-coupled agarose beads. The precipitated proteins were analyzed by immunoblotting with anti-GFP and anti-FLAG antibodies. **(I)** Co-immunoprecipitation assay of the interaction between TGA6 and MED15 in transgenic *Arabidopsis* expressing TGA6-GFP and MED15-HA-TurboID or mCherry-HA-TurboID. Immunoprecipitation was conducted using anti-HA magnetic beads, followed by immunoblotting with anti-GFP and anti-HA antibodies. The experiments in (**A**-**C** and **E**-**F**) were repeated three times and in (**H**-**I**) were repeated twice with similar results.

To further validate this interaction, we performed proximity labeling and co-IP assays. NPR1-GFP was robustly biotinylated when co-expressed with MED15-HA-TurboID, but not with LBD16-HA-TurboID in transgenic *Arabidopsis* plants (Figure 5E). LBD16, a transcription factor, was included as a negative control. Moreover, NPR1-GFP was co-IPed with MED15-HA-TurboID (Figure 5F). Upon NADP^+^ treatment, the NPR1-GFP/MED15-HA-TurboID association exhibited a biphasic pattern, with a gradual increase in association up to 8 hr post-treatment, followed by a decline to below basal levels by 24 hr (Figure 5F). These findings further support the *in vivo* association between NPR1 and MED15.

Since NPR1 interacts with MED15, we next examined whether TGAs also interact with MED15. Y2H assays showed that yeast cells expressing MED15 as bait and TGA1, TGA2, TGA5, TGA6, or TGA7 as prey grew on TDO medium supplemented with 5 mM 3-AT, whereas those expressing TGA3 or TGA4 as prey did not (Figure 5G). In contrast, yeast cells expressing MED14 or MED16 as bait with any TGA as prey did not grow (Supplemental Figure 8A), suggesting that MED15 is the primary Mediator subunit interacting with TGAs. To confirm this interaction *in vivo*, we conducted co-IP assays. When transiently co-expressed in *N. benthamiana*, TGA2-GFP, TGA5-GFP, and TGA6-GFP, but not GFP alone, were co-IPed with MED15-FLAG (Figure 5H; Supplemental Figures 8B and 8C). Additionally, TGA6-GFP was co-IPed with MED15-HA-TurboID, but not with mCherry-HA-TurboID, from transgenic *Arabidopsis* plants (Figure 5I), further validating the interaction between MED15 and TGAs.

### LecRK-VI.2-mediated NPR1 phosphorylation enhances TGA-MED15 interaction

Given that NPR1 interacts with both TGAs and MED15, we investigated whether NPR1 modulates TGA-MED15 association. In *N. benthamiana*, co-IP assays showed that TGA6-GFP associated with MED15-FLAG, and this interaction was markedly enhanced upon co-expression of NPR1-MYC (Figure 6A), suggesting that NPR1 promotes TGA6-MED15 complex formation. Consistently, in transgenic *Arabidopsis*, TGA6-GFP was weakly co-IPed with MED15-HA-TurboID in *npr1-3* plants (Figure 6B lane 1), indicating that a basal interaction can occur independently of NPR1. This association was strengthened in *npr1-3* plants complemented with *NPR1-GFP* (lane 2), Notably, NADP^+^ treatment did not increase the TGA6-GFP-MED15 interaction in the absence of NPR1-GFP (lane 3), but significantly enhanced it in the presence of NPR1-GFP (lane 4). Furthermore, NPR1-GFP was co-IPed with both TGA6-GFP and MED15-HA-TurboID upon NADP^+^ treatment (lane 4), indicating the *in vivo* formation of a ternary TGA6-NPR1-MED15 complex. Together, these findings suggest a model in which NPR1 functionally bridges TGA and MED15 following LecRK activation, enhancing the assembly of a transcriptional complex required for eNAD(P)-responsive gene expression.

**Figure 6.**
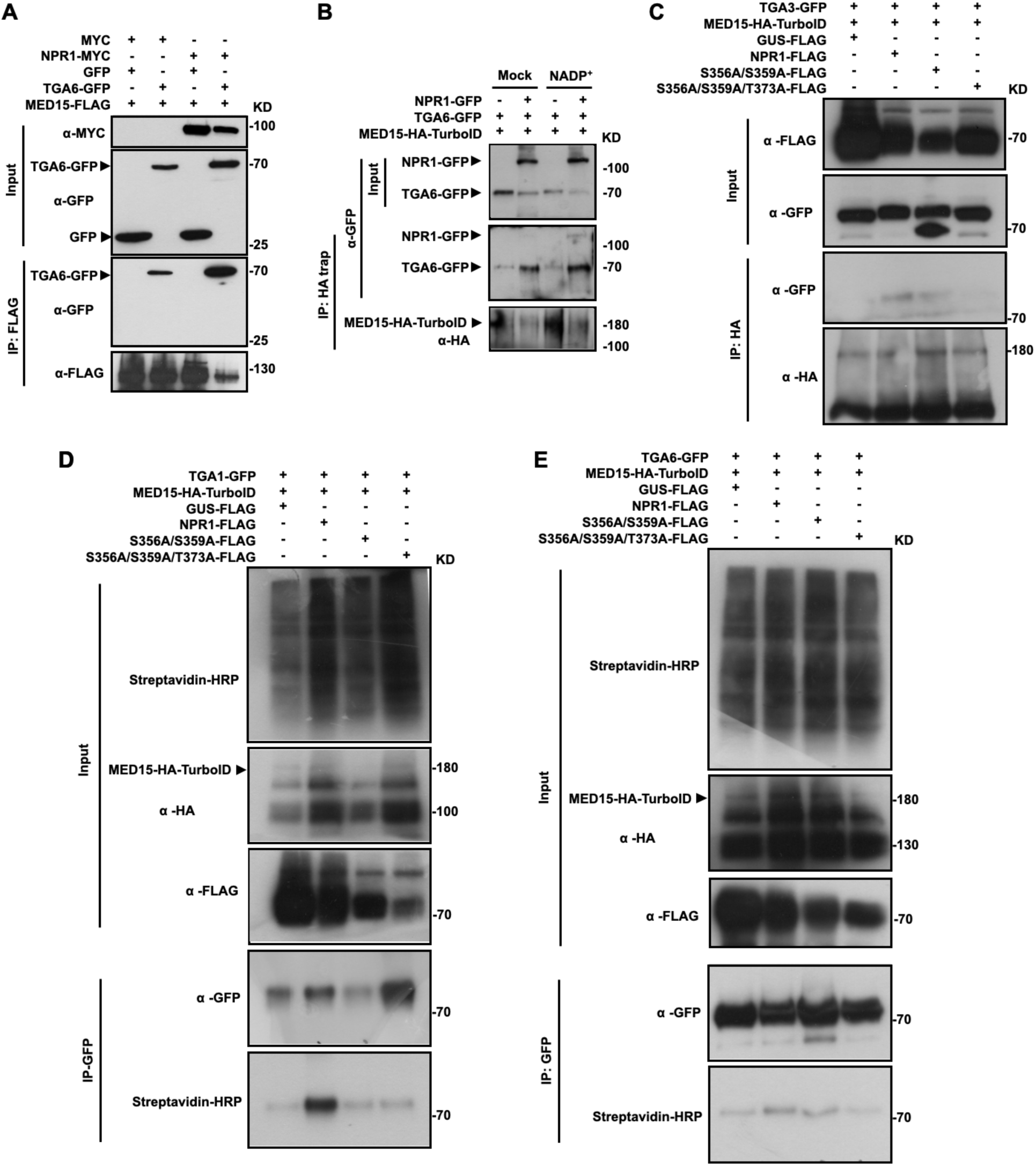
NPR1 phosphorylation by LecRK-VI.2 promotes TGA association with MED15. **(A)** Co-immunoprecipitation assay assessing the effect of NPR1 on the TGA6-MED15 interaction in *N. benthamiana*. The indicated proteins were transiently co-expressed in *N. benthamiana*. Immunoprecipitation was performed using anti-FLAG antibody-coupled agarose beads, followed by immunoblotting with anti-GFP and anti-FLAG antibodies. **(B)** Co-immunoprecipitation assay assessing the effect of NPR1 on the TGA6-MED15 interaction in *Arabidopsis* with or without NADP^+^ treatment. Leaves on transgenic *npr1-3* plants expressing TGA6-GFP and MED15-HA-TurboID, with or without NPR1-GFP, were treated with 0.8 mM NADP^+^ or water (mock) for six hours. Total protein was extracted and subjected to immunoprecipitation with anti-HA magnetic beads, followed by immunoblotting with anti-GFP and anti-HA antibodies. **(C)** Co-immunoprecipitation assay examining how nonphosphorylatable mutations affect NPR1-enhanced TGA3-MED15 interaction. NPR1-FLAG, S356A/T359A-FLAG, S356A/T359A/T373A-FLAG, or GUS-FLAG were transiently co-expressed with TGA3-GFP and MED15-HA-TurboID in *N. benthamiana*. Immunoprecipitation was carried out using anti-HA magnetic beads. The precipitated proteins were analyzed by immunoblotting with anti-GFP and anti-HA antibodies. **(D** and **E)** TurboID-based proximity labeling examining the effects of nonphosphorylatable mutations on NPR1-enhanced TGA1-MED15 (**D**) and TGA6-MED15 (**E**) association. Total protein was extracted from biotin-treated *N. benthamiana* leaves co-expressing the indicated proteins and subjected to immunoprecipitation with GFP-trap magnetic beads, followed by immunoblotting with streptavidin-HRP and anti-GFP antibody. The experiments in (**A**, **B**, and **E**) were repeated twice and in (**C** and **D**) were repeated three times with similar results.

To determine whether LecRK-VI.2-mediated phosphorylation of NPR1 is required for promoting TGA-MED15 association, we tested nonphosphorylatable NPR1 variants. T373, an *in vivo* phosphorylation site (Yao *et al*., 2023), was phosphorylated by LecRK-VI.2 *in vitro* (Supplemental Figure 2D). A T373A substitution impaired NPR1 function in NAD(P)^+^-induced immunity and SAR (Supplemental Figures 9A-9D) (Lee et al., 2015), weakened its interaction with TGA3 (Supplemental Figure 10A), and when combined with S356A and T359A, additively disrupted its association with MED15 in yeast (Supplemental Figure 10B). We therefore included an *S356A/T359A/T373A* triple mutant in our assays. Wild-type NPR1-FLAG significantly enhanced the amount of TGA3-GFP co-IPed with MED15-HA-TurboID, whereas *S356A/T359A-FLAG* had only a minor effect, and the triple mutant abolished the interaction (Figure 6C). Similarly, wild-type NPR1-FLAG markedly increased biotinylation of TGA1-GFP and TGA6-GFP co-expressed with MED15-HA-TurboID, while both *S356A/T359A-FLAG* and the triple mutant showed reduced or no effect (Figures 6D and 6E). These results demonstrate that LecRK-VI.2-mediated phosphorylation of NPR1 is essential for strengthening the TGA-MED15 interaction.

## Discussion

Since SAR was first described over six decades ago (Ross, 1961), defining its underlying signaling pathway has remained a central challenge in plant immunity (Durrant and Dong, 2004; Kachroo and Kachroo, 2020; Shah and Zeier, 2013; Vlot et al., 2021). Early advances established key components of SAR, including the identification of pathogenesis-related (PR) proteins in the 1970s (van Loon and van Kammen, 1970; Van Loon and Van Strien, 1999), the recognition of SA as a core signaling molecule in the 1990s (Malamy et al., 1990; Métraux et al., 1990), and the cloning of the master transcriptional coactivator NPR1 in 1997 (Cao *et al*., 1997; Ryals et al., 1997; Shah et al., 1997), later confirmed as an SA receptor in 2012 (Wu et al., 2012). Beginning in the early 2000s, multiple candidate mobile signals were proposed to mediate long-distance SAR (Chanda *et al*., 2011; Chaturvedi *et al*., 2008; Chen *et al*., 2018; Hartmann *et al*., 2018; Jung *et al*., 2009; Li *et al*., 2023; Maldonado *et al*., 2002; Navarova *et al*., 2012; Park *et al*., 2007; Riedlmeier *et al*., 2017; Shine *et al*., 2022; Shulaev *et al*., 1997; Wang *et al*., 2018), yet how these signals are perceived in systemic tissues and translated into transcriptional outputs has remained largely unresolved.

Our recent work identifies eNAD(P) as a critical integrative signal downstream of diverse SAR-associated cues and establishes the receptor kinase LecRK-VI.2 as its sensor in systemic tissues (Li *et al*., 2023; Wang *et al*., 2019). However, a fundamental mechanistic gap persists between signal perception at the cell surface and activation of NPR1-dependent transcription.

Here, we show that LecRK-VI.2 directly phosphorylates NPR1 (Figures 1 and 2; Supplemental Figure 2), enabling its productive interaction with TGAs and the Mediator complex to drive defense gene expression (Figure 4; Supplemental Figure 10B). These findings define a receptor-to-coactivator signaling mechanism that directly links extracellular signal perception to transcriptional activation, thereby closing a major gap in the SAR signaling pathway and establishing a coherent framework from signal perception to transcriptional reprogramming in systemic immunity (Figure 7).

**Figure 7.**
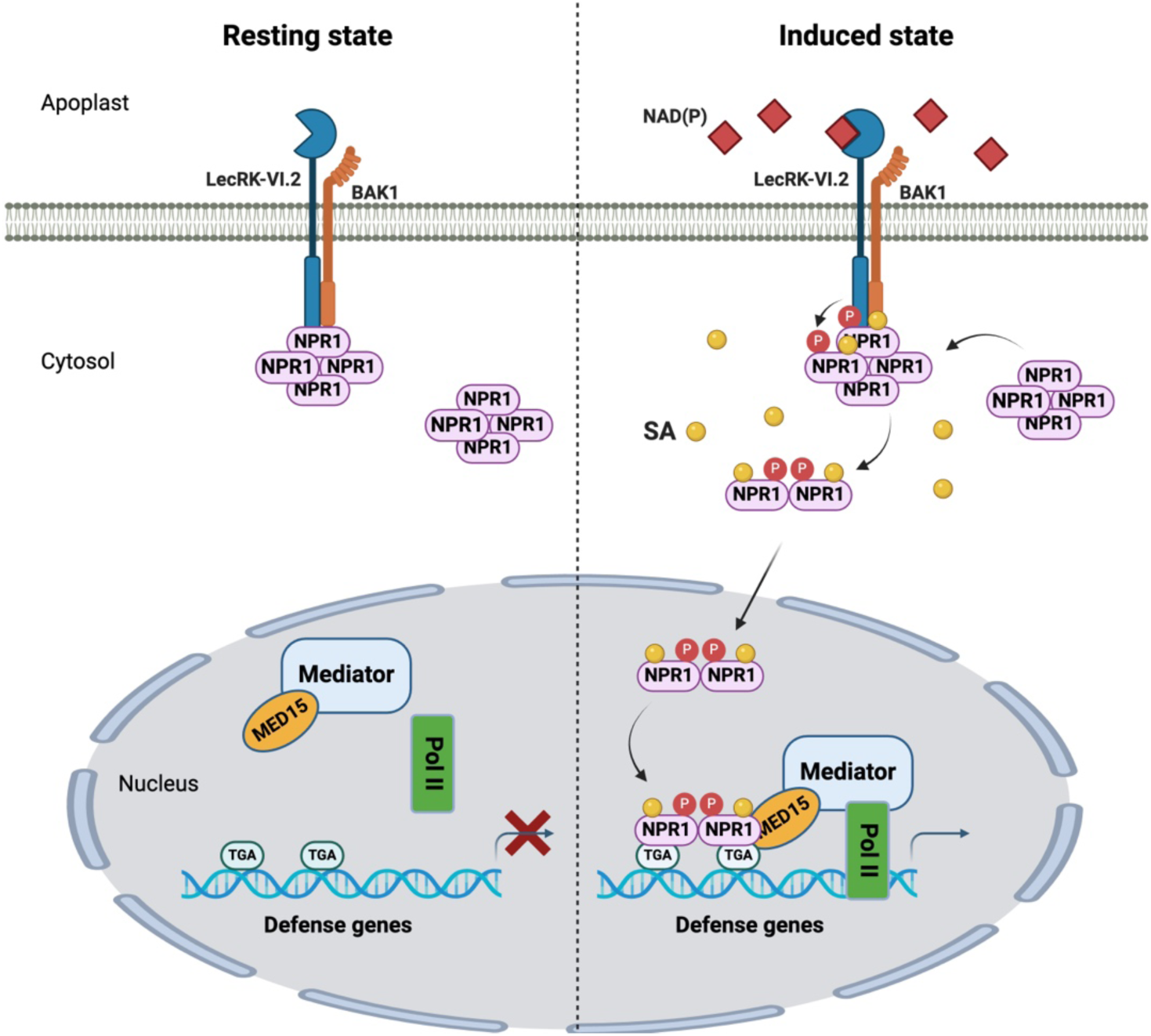
Proposed model illustrating how LecRK-VI.2 and NPR1 mediate eNAD(P) signal transduction. In the resting state, a subset of NPR1 oligomers associates with the eNAD(P) receptor complex LecRK-VI.2/BAK1. Upon pathogen infection, NAD(P) is released into the extracellular space, where it binds LecRK-VI.2, triggering receptor-mediated phosphorylation of NPR1 at S356, T359, and T373. This modification is followed by dissociation of NPR1 from LecRK-VI.2. Extracellular NAD(P) also promotes SA accumulation through unknown mechanisms (Zhang and Mou, 2009), altering the cellular redox state and shifting NPR1 from oligomers to dimers (Mou *et al*., 2003). SA binding to NPR1 facilitates docking of its SA-binding domain onto ankyrin repeats 3 and 4 (Kumar *et al*., 2022), a key conformational change that primes NPR1 for transcriptional activation. In the nucleus, phosphorylation at S356, T359, and T373 enhances NPR1 interaction with TGAs and the Mediator subunit MED15, promoting assembly of the transcription initiation complex and activation of immune-related gene expression. The figure was created using the BioRender online tool (https://www.biorender.com/).

NPR1 functions as a central transcriptional coactivator that integrates diverse immune signals to activate defense gene expression, primarily through interaction with TGAs and other transcription regulators (Chen et al., 2019; Després *et al*., 2000; Powers *et al*., 2024; Zavaliev and Dong, 2024; Zhang *et al*., 1999; Zhou *et al*., 2000). Consistent with this role, NPR1 associates with multiple chromatin and transcriptional regulators, including histone modifiers and Mediator subunits (Chen *et al*., 2019; Jin et al., 2018; Powers *et al*., 2024). Our data identify MED15 as a key NPR1-interacting partner and demonstrate that NPR1 enhances TGA-MED15 association (Figures 5D-5F, 6A and 6B), supporting assembly of an active transcriptional complex. These findings position NPR1 as a central hub linking extracellular signal perception to the core transcriptional machinery.

Importantly, we show that LecRK-VI.2-mediated phosphorylation is required for NPR1 interactions with both TGAs and MED15 (Figures 4A-4G; Supplemental Figure 10B), as well as for the enhanced association between TGAs and MED15 (Figures 6C-6E). Nonphosphorylatable NPR1 variants disrupt these interactions and abolish SAR (Figures 3C, 3F and 3G), indicating that this modification functions as a molecular switch controlling NPR1 activity. We propose that phosphorylation enables NPR1 to transition into a transcriptionally competent state, enabling efficient assembly of a higher-order transcriptional complex (Rochon *et al*., 2006; Zavaliev et al., 2020). Given the breadth of the NPR1 interactome (Powers *et al*., 2024), this modification is likely to have widespread effects on NPR1-dependent transcriptional programs.

Our results further define a temporal sequence in which LecRK-VI.2-mediated phosphorylation occurs rapidly upon eNAD(P) perception and precedes NPR1 engagement with transcriptional partners (Figure 2F; Supplemental Figures 6B-6D). However, phosphorylation alone is insufficient for full NPR1 function, as phosphomimetic NPR1 variants do not exhibit constitutive activity (Figures 3A-3E). This suggests that additional regulatory inputs are required. Because eNAD(P)-induced immunity depends on SA (Wang *et al*., 2019; Zhang and Mou, 2009), and SA regulates NPR1 through multiple post-translational mechanisms, including S55/S59 dephosphorylation, SUMOylation, S11/S15 phosphorylation, and ubiquitination (Mou et al., 2003; Saleh et al., 2015; Spoel et al., 2009), it is likely that these pathways act in concert. In this context, LecRK-VI.2-mediated phosphorylation may function as a priming event that licenses NPR1 for subsequent SA-dependent activation, thereby ensuring temporal coordination and signaling specificity.

Cell-surface immune receptors, particularly receptor kinases, typically transmit signals through multi-step phosphorylation cascades involving receptor-like cytoplasmic kinases and MAP kinases to regulate transcriptional outputs (Bender and Zipfel, 2023). In some cases, receptor kinases can directly phosphorylate transcription factors (David et al., 1996; Macias-Silva et al., 1996), providing a more direct route to gene regulation. However, such mechanisms remain relatively rare in plants (Guo et al., 2018; Meyer et al., 2015), and direct targeting of a transcriptional coactivator by a cell-surface receptor kinase has not been demonstrated in plants and remains rarely described more broadly. Here, we uncover a receptor-coactivator signaling module in which LecRK-VI.2 directly phosphorylates NPR1 to couple eNAD(P) perception to transcriptional activation. This mechanism establishes a direct, receptor-proximal route from cell-surface signal perception to coactivator-driven gene expression, bypassing canonical multi-step cascades. By linking extracellular immune cues to the core transcriptional machinery, our findings define a fundamental principle for signal integration in systemic immunity and reveal receptor-to-coactivator coupling as a mechanism for rapid conversion of extracellular signals into coordinated transcriptional outputs (Figure 7).

## Methods

### Arabidopsis

*Arabidopsis thaliana* mutants and transgenic lines used in this study were all in the Col-0 ecotype background. Transgenic *Arabidopsis* expressing GFP-fused NPR1, OsNPR1, BrNPR1, or their mutant variants are in the *npr1-3* background. Transgenic *Arabidopsis* expressing NPR1-GFP in *lecrk-VI.2-1* background were generated by crossing *35S:NPR1-GFP/npr1-3* with the *lecrk-VI.2-1* mutant. Double transgenic *Arabidopsis* lines expressing NPR1-GFP and LecRK-VI.2-HA-TurboID or Lti6b-HA-TurboID were generated by dipping the *35S:NPR1-GFP/npr1-3* transgenic plants with the *Agrobacterium tumefaciens* strain EHA105 harboring the *35S:LecRK-VI.2-HA-TurboID* or *35S:Lti6b-HA-TurboID* constructs, respectively. Double transgenic *Arabidopsis* lines expressing NPR1-GFP, or its phospho-mutant variants and TGAs-HA-TurboID were generated by dipping the *35S:NPR1-GFP/npr1-3*, *35S:npr1^S356A/T359A^-GFP/npr1-3*, or *35S:npr1^S356D/T359D^-GFP/npr1-3* transgenic plants with the *A. tumefaciens* strain EHA105 harboring the *35S:TGAs-HA-TurboID* constructs. Double transgenic *Arabidopsis* lines expressing TGA6-GFP and MED15-HA-TurboID or mCherry-HA-TurboID were generated by dipping the *35S:TGA6-GFP* transgenic plants with the *A. tumefaciens* strain EHA105 harboring the *35S:MED15-HA-TurboID* or *35S:mCherry-HA-TurboID* constructs, respectively.

*Arabidopsis* lines expressing NPR1-GFP and MED15-HA-TurboID or LBD16-HA-TurboID were generated by dipping the *35S:NPR1-GFP* transgenic plants with the *A. tumefaciens* strain EHA105 harboring the *35S:MED15-HA-TurboID* or *35S:LBD16-HA-TurboID* construct, respectively. Double transgenic *Arabidopsis* expressing TGA6-GFP and MED15-HA-TurboID in *npr1-3* background were generated by dipping the *35S:MED15-HA-TurboID/npr1-3* transgenic plants with the *A. tumefaciens* strain EHA105 harboring *35S:TGA6-GFP*. Triple transgenic *Arabidopsis* expressing TGA6-GFP, MED15-HA-TurboID and NPR1-GFP in *npr1-3* background were generated by dipping the *35S:MED15-HA-TurboID/35S:NPR1-GFP/npr1-3* plants with the *A. tumefaciens* strain EHA105 carrying *35S:TGA6-GFP*. *Arabidopsis* and *Nicotiana benthamiana* seeds were stratified at 4°C for three days before germination, and all the plants for experiments were grown in soil in a growth room at 24°C/22°C (day/night) and 60% relative humidity with a 14 hr/10 hr light/dark photoperiod. For selection of transgenic *Arabidopsis* plants with antibiotic resistance, seeds were surface sterilized with 85% (v/v) ethanol, dried on filter paper, and germinated on sterile half-strength (½) MS medium (pH 5.7) supplemented with 1% (w/v) sucrose and 0.55% (w/v) agar with appropriate antibiotics.

### Plasmid construction

The coding sequences of the *Arabidopsis* genes *NPR1*, *BAK1*, *LecRK-VI.2*, *LecRK-I.8*, *Lti6b*, and *TGAs* were amplified from the cDNA of Col-0. To generate transgenic *Arabidopsis* lines expressing NPR1-GFP, the coding sequence of *NPR1* was amplified and inserted into SacI/SalI-linearized pCAMBIA1300S-GFP (Wang *et al*., 2019) using In-Fusion Snap Assembly (Takara). To generate transgenic *Arabidopsis* lines expressing BrNPR1-GFP, the coding sequence of *BrNPR1* was amplified from the cDNA of *B. rapa* var. R500 and inserted into KpnI/BamHI-digested pCAMBIA1300S-GFP using In-Fusion Snap Assembly (Takara). To generate transgenic *Arabidopsis* lines expressing OsNPR1-GFP, codon-optimized *OsNPR1* coding sequence was synthesized and inserted into KpnI/BamHI-digested pCAMBIA1300S-GFP using In-Fusion Snap Assembly (Takara). To express LecRK-VI.2-HA-TurboID and Lti6b-HA-TurboID, the coding sequences of *LecRK-VI.2* and *Lti6b* were amplified and inserted into XhoI/XbaI-linearized pBASTA-HA-TurboID (Wu et al., 2020) using T4 ligase. To express TGAs-HA-TurboID, the coding sequences of *TGA1*, *TGA2*, and *TGA3* were amplified and inserted into KpnI/XhoI-linearized pBASTA-HA-TurboID using T4 ligase. To express MED15-HA-TurboID, LBD16-HA-TurboID, and mCherry-HA-TurboID, the coding sequences of *MED15*, *LBD16*, and *mCherry* were amplified and inserted into KpnI/BamHI-linearized pBASTA-HA-TurboID using T4 ligase. For transient expression in *N. benthamiana*, the coding sequences of *TGA1*, *TGA2*, *TGA3*, *TGA5*, and *TGA6* were amplified and inserted into BamHI/SalI-digested pCAMBIA1300S-GFP, and those of *NPR1*, *MED15*, and *GUS* were amplified and inserted into SacI/SalI-digested pCAMBIA1300S-FLAG (Wang et al., 2015), respectively, using T4 ligase or In-Fusion Snap Assembly (Takara). For yeast two-hybrid assays, the fragments encoding the kinase domains (KDs) of LecRK-VI.2KD, BAK1KD, and LecRK-I.8KD as well as the full-length coding sequences of MED14, MED15, MED16 were amplified and inserted into pGBKT7, and the coding sequence of *NPR1* was inserted into pGADT7 using In-Fusion Snap Assembly (Takara). The bait and prey vectors were linearized by NdeI and BamHI digestion before the In-Fusion reactions. For *in vitro* kinase assay, the coding sequence of *NPR1* was amplified and inserted into pET28a-sfGFP-His using Golden Gate Assembly. pMAL-p2X-MBP-LecRK-VI.2KD and pGEX-4T-1-GST-BAK1KD have been described previously (Wang *et al*., 2019). For SLC assays, the coding sequence of *NPR1* was inserted into SacI/BamHI digested pCAMBIA1300S-NlucC-FLAG using In-Fusion Snap Assembly. pCAMBIA1300S-LecRK-VI.2-NlucN-HA and pCAMBIA1300S-LecRK-I.8-NlucN-HA have been described previously (Li *et al*., 2024). Point mutations in *NPR1*, *OsNPR1*, *BrNPR1*, and *LecRK-VI.2* were generated using the QuikChange site-directed mutagenesis kit (Agilent) or overlap extension PCR. All the primers used for plasmid construction were listed in Supplemental Table 1.

### *In vitro* kinase assay

The *in vitro* kinase assay was performed as described previously (Liu et al., 2019) with slight modifications. Briefly, 5 μg of indicated recombinant proteins were incubated in 30 μL kinase assay buffer (25 mM Tris-HCl, pH 7.5, 10 mM MnCl_2_, 10 mM MgCl_2_, 10 mM ATP) at 30°C for 1 hr. The reaction was stopped by adding 8 mL of Laemmli buffer, and the sample was boiled for 8 min and separated in 8% SDS-PAGE gel. Phosphorylation of NPR1 was detected by immunoblotting with an anti-pThr antibody (1:1000, Cell signaling).

### LC-MS/MS detection of NPR1 phosphorylation sites

To identify *in vitro* LecRK-VI.2 phosphorylation sites on NPR1, purified recombinant proteins sfGFP-NPR1-His and MBP-LecRK-VI.2KD or GST-BAK1KD were incubated in kinase assay buffer (25 mM Tris-HCl, pH 7.5, 10 mM MnCl_2_, 10 mM MgCl_2_, 10mM ATP) at 30°C for 1 h with gentle shaking. The reaction was terminated by adding Laemmli buffer, and the samples were boiled for 8 min and separated in 8% SDS-PAGE gel. The SDS-PAGE gel was then stained with Coomassie Brilliant Blue (CBB) R250, and the sfGFP-NPR1-His bands were cut out for LC-MS/MS analysis. Protein in-gel trypsin-digestion, peptides extraction with acetonitrile, and phosphopeptides enrichment using a NuTip TiO_2_ + ZrO_2_ (Glygen Inc. NT2TIZR) were described in detail by Perron et al (Perron et al., 2022; Ye et al., 2022). Ten μL of acidified phospho-enriched peptide sample was injected on Acclaim PepMap 100 C18 precolumn (20 mm by 75 μm; 3 μm) and separated on a PepMap RSLC C18 analytical column (250 mm by 75 μm; 2 μm) at a flow rate of 250 nl/min followed by LC-MS/MS on Easy-nLC and Orbitrap Fusion mass spectrometer (Thermo Fisher Scientific). Mobile phase A was water with 0.1% formic acid and mobile phase B was acetonitrile with 0.1% formic acid. A 90 min gradient was used for separation in positive mode. The gradient started at 2% B, increased to 5% B till 5 min, linearly increased to 40% till 70 min, 45% till 75 min, and ended at 98% B at 90 min. A collision-induced dissociation (CID) and electron transfer dissociation (ETD) decision tree method was used for this experiment. The Orbitrap MS1 resolution was 120 k, the scan range was 350 to 1800 m/z, the nanoelectrospray voltage was 2000, the automatic gain control target was set to 200,000, microscan was 1, the RF lens was 55% and the maximum inject time was set to 50 ms. The MS/MS spectra were acquired in the linear ion trap with ETD (with charge states 6-8) and CID (with charge state 2-5), a mass window of 1.3, collision energy of 35, activation time of 10 ms, and activation q of 0.25. To identify NADP^+^-induced phosphorylation sites on NPR1 *in vivo*, leaves of four-week-old double transgenic plants expressing NPR1-GFP and LecRK-VI.2-HA-TurboID were infiltrated with 0.8 mM NADP^+^ or water. Leaf tissues were collected 30 min later. Total proteins were extracted from 20 g collected leaf tissues with extraction buffer (50 mM HEPES-KOH, pH 7.5, 150 mM KCl, 1 mM EDTA, 0.5% Triton X-100, 5 mM NaF, 5 mM Na_3_VO_4_, and proteinase inhibitor cocktail). To precipitate NPR1-GFP, the total proteins were incubated with precleared protein A agarose beads (Santa Cruz) and the anti-NPR1 antibody for 3 hr at 4°C with rotation. After immunoprecipitation, the beads were washed three times with extraction buffer without proteinase inhibitor cocktail and boiled in Laemmli buffer for 8 min to elute the bound proteins. The eluted proteins were separated in 8% SDS-PAGE gel and stained with CBB. The NPR1-GFP band was cut for trypsin digestion and LC-MS/MS analysis as described above.

### Phospho-specific antibody development

Rabbit polyclonal anti-pT359 antibody was developed by Pierce/Thermo Fisher Scientific using an NPR1-pT359 peptide [C-ASASEA(pT)LEGRTA-amide]. The specificity of the newly developed anti-pT359 antibody was confirmed using LecRK-VI.2KD-treated NPR1 and npr1^T359A^ (Figure 2E). The anti-pT359 antibody was diluted 1:2000 for detecting NADP^+^- and SA-induced phosphorylation of T359 *in vivo* and for phosphorylation of T359 by LecRK-VI.2KD in *in vitro* kinase assays.

### Phos-tag gel assay

Phos-tag gel assays of *in vitro* NPR1 phosphorylation were conducted according to the Phos-tag™ SDS-PAGE guidebook (FUJIFILM wako). Briefly, the recombinant proteins sfGFP-ANK-His or the corresponding nonphosphorylatable variants were incubated with MBP-LecRK-VI.2KD or the kinase dead version MBP-LecRK-VI.2KD^D494N^ in 30 μL kinase assay buffer (25 mM Tris-HCl, pH 7.5, 10 mM MnCl_2_, 10 mM MgCl_2_, 10 mM ATP) at 30°C for 1 hr. The reaction was stopped by adding 8 uL of Laemmli buffer, and the sample was boiled for 8 min and separated in 10 % SDS-PAGE gel, and in a 6 % SDS-PAGE gel containing 50 μM Phos-tag acrylamide AAL-107 (NARD Institute, FUJIFILM wako) and 100 μM MnCl_2_. After 2 h electrophoresis at 120V, the phos-tag gel was treated with transfer buffer containing 10 mM EDTA for three times (each time for 10 min) and then soaked in transfer buffer without EDTA twice (each time for 10 min). The proteins were transferred to a polyvinylidene fluoride (PVDF) membrane using transfer buffer at 25V overnight. The membrane from the phos-tag gel was blocked with 5% fat-free milk and probed with anti-GFP antibody (1:5,000, Clontech). The membrane from the regular SDS-PAGE gel was blotted with anti-GFP antibody (1:5,000, Clontech), anti-MBP antibody (1:10,000, NEB), and anti-pThr antibody (1:1000, Cell signaling).

Phos-tag gel assays of *in vivo* NPR1 phosphorylation were conducted according to the Phos-tag™ SDS-PAGE guidebook (FUJIFILM wako). Briefly, four-week-old plants were treated with 0.8 mM NADP^+^ or water for 45 min, ground in liquid nitrogen, and homogenized in modified extraction buffer with a low concentration of salt (25 mM Tris-HCl, pH 7.5, 10 mM NaCl, 5 mM NaF, 5 mM Na_3_VO_4_, and EDTA-free protease inhibitor cocktail). The lysate was centrifuged at 16000 g for 15 min at 4°C, and the supernatant was collected. The extracted proteins were resolved in a 6% SDS-PAGE gel containing 50 μM Phos-tag acrylamide AAL-107 (NARD Institute, FUJIFILM wako) and 100 μM ZnCl_2_. After 6 h electrophoresis at 120V, the gel was treated with transfer buffer containing 10 mM EDTA for three times (each time for 10 min) and then soaked in transfer buffer without EDTA twice (each time for 10 min). The proteins were transferred to a polyvinylidene fluoride (PVDF) membrane using transfer buffer at 25V overnight. The membrane was blocked with 5% fat-free milk and probed with anti-GFP antibody (1:5,000, Clontech).

### Yeast two-hybrid assay

The yeast two-hybrid assays were performed according to the Matchmaker GAL4 Two-Hybrid System manual (Clontech). Briefly, paired bait and prey constructs were co-transformed into the yeast strain AH109, and the resulting cells were plated on synthetic dextrose (SD) double dropout (DDO) (-Trp-Leu) agar plates. Yeast colonies formed on the DDO medium were suspended in sterile water and then simultaneously dropped onto DDO and selective agar plates SD triple dropout (TDO) (-Trp-Leu-His) + 5 mM 3-amino-1,2,4-triazole (3-AT) or SD quadruple dropout (QDO) (-Trp-Leu-His-Ade) medium. The protein-protein interactions were determined by the growth rate of yeast cells on the selective medium.

### Split Nano luciferase complementation (SLC) assay

SLC assays were performed as previously described (Wang *et al*., 2020). Briefly, different combinations of *A. tumefaciens* strain EHA105 cells harboring the indicated constructs were co-infiltrated into fully expanded leaves of five-week-old *N. benthamiana* plants. Three days later, 20 μM Coelenterazine h (Nanolight 301) were applied to the infiltrated leaves by infiltration.

Leaf disks (7 mm in diameter) were immediately collected with a hole punch, and each leaf disc was settled in a well of a 96-well white plate (150 μL water/well). The luciferase activities were measured using a GloMax Discover luminometer (Promega). After measurement, the leaf disks were kept for protein abundance analysis. The indicated NlucN/NlucC fusion proteins were detected by immunoblotting with anti-HA (1:3000, Roche) and anti-FLAG (1:1000, Sigma) antibodies.

### Co-immunoprecipitation (co-IP) assay

Co-IP assays in *N. benthamiana* were performed as previously described (Liu *et al*., 2019). Briefly, designated genes were transiently expressed in *N. benthamiana* leaves, and total proteins were extracted using an extraction buffer containing 50 mM HEPES-KOH (pH 7.5), 150 mM KCl, 1 mM EDTA, 0.5% Triton X-100, and proteinase inhibitor cocktail. For IP of GFP-fusion proteins, extracts were incubated for 2 hr at 4°C with 6 μL of GFP-trap magnetic agarose beads (Chromotek), followed by five washes with wash buffer (50 mM HEPES-KOH, pH 7.5, 150 mM KCl, 1 mM EDTA, 0.5% Triton X-100). Precipitated proteins were eluted by adding 60 μL of Laemmli buffer and boiling for 8 min. For IP of FLAG-fusion proteins, total proteins were incubated for 2 hr at 4°C with 8 μL of anti-FLAG antibody-coupled agarose beads (Sigma), followed by six washes with wash buffer. Bound proteins were eluted using 100 μL 3 × FLAG peptides (Sigma). The precipitated proteins were separated in 8% SDS-PAGE gels and analyzed by immunoblotting with anti-GFP (1:5000, Clontech) and anti-FLAG (1:1000, Sigma) antibodies. For time-course co-IP assays, leaves of double transgenic *Arabidopsis* plants were infiltrated with 0.8 mM NADP^+^, collected at the indicated time points, and snap-frozen in liquid nitrogen. Total proteins were extracted with extraction buffer and incubated for 2 hr at 4°C with 6 μL of anti-HA antibody-coupled magnetic agarose beads (Cell signaling). Beads were washed six times with wash buffer, and precipitated proteins were eluted by boiling for 8 min in 60 μL of Laemmli buffer, separated in 8% SDS-PAGE gels, and analyzed by immunoblotting with anti-GFP (1:5000, Clontech) and anti-HA (1:3000, Roche) antibodies.

### TurboID-based proximity labeling assay

TurboID-based proximity labeling assays were performed as described previously (Wu *et al*., 2020). Briefly, leaves on *Arabidopsis 35S:LecRK-VI.2-HA-TurboID/35S:NPR1-GFP* and *35S:Lti6b-HA-TurboID/35S:NPR1-GFP* double transgenic plants were infiltrated with 50 μM biotin and incubated at room temperature for 3 hr to allow labeling. For TurboID-based proximity labeling assays in *N. benthamiana,* different combinations of *A. tumefaciens* strain EHA105 cells harboring the indicated constructs were co-infiltrated into fully expanded leaves of five-week-old *N. benthamiana* plants. Two days later, the infiltrated leaves were treated with 200 uM biotin and incubated at room temperature for 3 hr to allow labeling. Total proteins were extracted using an extraction buffer containing 50 mM HEPES-KOH (pH 7.5), 150 mM KCl, 1 mM EDTA, 0.5% Triton X-100, and proteinase inhibitor cocktail. To enrich NPR1-GFP, the extracts were incubated with GFP-trap Magnetic Agarose beads (ChromoTek) for 2 hr at 4°C. The beads were then washed three times with the extraction buffer, omitting the proteinase inhibitors. The bound proteins were eluted by adding Laemmli buffer and boiling for 8 min. Biotinylation was detected using Streptavidin-HRP (1:20000, Abcam).

### Promoter transactivation assay

The *PR1* promoter reporter (*pPR1:DUAL-LUC*) was co-infiltrated with *35:GFP*, *35S:NPR1-GFP*, *35S:npr1^S356A/T359A^-GFP*, or *35S:npr1^S356D/T359D^-GFP* into *N. benthamiana* leaves. Two days post-infiltration, 7 mm leaf disks were collected from the infiltrated leaves, ground in liquid nitrogen, and lysed with the PLB buffer from the Dual-Luciferase Reporter Assay System (Promega E1910) for 15 min with gentle rocking at room temperature. The lysate was centrifuged at top speed for 1 min, and 20 μL of the supernatant was used to measure FLUC and RLUC activities according to the manufacturer’s instructions, using a GloMax Discover luminometer (Promega). Briefly, at 25°C, substrates for FLUC and RLUC were added using an automatic injector. Following a 2-s premeasurement delay, signals were captured for 10 s and recorded as counts per second. *PR1* promoter activity was determined by calculating the ratio of F-LUC and R-LUC activities for each effector and plotting it relative to free GFP, which was arbitrarily set to 1.

### RNA isolation and quantitative PCR (qPCR)

Approximately 100 mg leaf tissues were snap-frozen in liquid nitrogen and ground into a fine powder in a 2 mL lysing tube with a Spex SamplePrep 2000 Geno/Grinder (OPS Diagnostics). Total mRNA was extracted using the E.Z.N.A.® Total RNA Kit (OMEGA), and reverse transcription was conducted using the SuperScript™ IV First-Strand Synthesis kit (Invitrogen), following the manufacturers’ manuals. qPCR was performed using SYBR™ Green PCR Master Mix (Applied Biosystems) on a QuantStudio 3 Real-Time PCR system (Applied Biosystems) according to the user’s manual. The 2^−ΔCt^ method was used to determine the relative level of gene expression. *UBQ5* was used as an internal control. The gene-specific primers used for qPCR were listed in Supplemental Table 1.

### NADP^+^-induced immunity and biological SAR assays

Four-week-old *Arabidopsis* plants were used for NADP^+^-induced immunity and biological SAR assays. The previously described pathogen strain *Pseudomonas syringae* pv*. maculicola* ES4326 with an integrated *luxCDABE* luciferase operon (*Psm_lux*) was used in this study (Li *et al*., 2023). *Psm_lux* was cultured at 28°C in King’s B medium [2% (w/v) proteose peptone, 0.15% (w/v) K_2_HPO_4_, 6 mM MgSO_4_, and 1.5% (v/v) glycerol] containing appropriate antibiotics. *Psm_lux* in overnight log-phase cultures were centrifuged at 5,000 rpm for 1 min to pellet cells. The pellet was resuspended in 5 mM sterile MgCl_2_ solution and diluted for leaf infiltration. NADP^+^-induced local and systemic immunity assays were performed as previously described with slight modifications (Li *et al*., 2023). For NADP^+^-induced local immunity, 0.8 mM freshly made NADP^+^ solution (pH = 5.7) or sterile water (negative control) was syringe-infiltrated into the 5^th^ and 6^th^ leaves from the bottom on each plant. After 4 hr, the NADP^+^-treated leaves were infiltrated with *Psm_lux* (OD_600_ = 0.001). *Psm_lux* growth in the leaves was determined 60 hr post inoculation (hpi). For NADP^+^-induced systemic immunity, 0.8 mM freshly made NADP^+^ solution (pH = 5.7) or sterile water was infiltrated into three lower leaves on each plant (the 3^rd^, 4^th^, and 5^th^ from the bottom). After 4 hr, two upper systemic leaves on the plant (the 6^th^ and 7^th^ from the bottom) were infiltrated with *Psm_lux* (OD_600_ = 0.001). *Psm_lux* growth in the leaves was assessed 60 hpi. Biological induction of SAR was conducted as previously described with slight modifications (Wang *et al*., 2019). Briefly, three lower leaves (the 3^rd^, 4^th^, and 5^th^ from bottom) were infiltrated with 5 mM MgCl_2_ (-SAR) or *Psm* (OD_600_ = 0.002) (+SAR). After 48 hr, two upper systemic leaves (the 6^th^ and 7^th^ from the bottom) were infiltrated with *Psm_lux* (OD_600_ = 0.001). *Psm_lux* growth in the leaves was assessed 60 hpi. For *Psm_lux* growth quantification, leaf disks (7 mm in diameter) were collected using a hole punch and placed in a white, light-reflecting 96-well plate (Corning 3912). One leaf disc was taken from each leaf. Leaf disks were floated on 150 μL 1 mM MgCl_2_ in each well to keep them wet. The plate containing leaf disks was placed in the sample drawer of a GloMax Discover luminometer (Promega) and kept in the dark by closing the lid for 10 min to reduce background signals. The relative light unit (RLU) of each sample was then measured for 10 s. Bacterial titers were expressed as log_10_(RLU) per leaf disc.

### Quantification and statistical analysis

The pathogen growth results in this study are based on one experimental dataset, confirmed through at least three independent experiments conducted at different times. For quantification of *Psm-lux* titers in the leaves, 8-12 *Arabidopsis* leaf replicates were used, with one leaf taken from each plant. In the SLC assays, six leaf disks were collected from treated *N. benthamiana* leaves and analyzed independently; the data were presented accordingly. This experiment was repeated three times, yielding similar results. For promoter transactivation assays, four leaf disks from four infiltrated *N. benthamiana* leaves were lysed together, and the lysate was analyzed four times, with data from the four technical replicates presented. The results were confirmed in three independent experiments conducted at different times. The gene expression results were derived from analyses of three independent biological samples, each taken from six *Arabidopsis* leaves on six plants (one leaf per plant). Statistical analyses were performed using Microsoft Excel (Student’s t-test) and Prism 10 (one-way/two-way ANOVA) on Microsoft Office 2023 for Macintosh.

## Data Availability

The authors declare that all data supporting the findings of this study are available within the manuscript and its supplemental files or are available from the corresponding author upon request. Source data are provided with this paper. This study did not generate large datasets.

## Author contributions

C.L., Q.Liu., and Z.M. conceived the project. C.L., Q.L., S. Chhajed, M.Z., F.M.H., Q.Li, X.Z., and S. Chen performed the experiments, analyzed the data, and prepared figures. C.L. and Z.M. wrote the manuscript with contributions from S. Chhajed and S. Chen. All authors provided comments for the manuscript before submission.

## Acknowledgments

We thank Dr. Yuelin Zhang at University of British Columbia for sharing with us the pBASTA-HA-TurboID vector and Dr. Xinnian Dong at Duke University for the pPR1:DUAL-LUC vector and the *npr1-3* and *tag2/3/5/6* mutant seeds. We are grateful to Dr. Pablo Tornero from Universidad Politécnica de Valencia-Consejo Superior de Investigaciones Científicas for sharing the *nrb4-3* seeds.

