## Supplemental figures and table for "Extracellular NAD(P) activates systemic acquired resistance through LecRK-VI.2-mediated phosphorylation of NPR1"

### Supplemental information

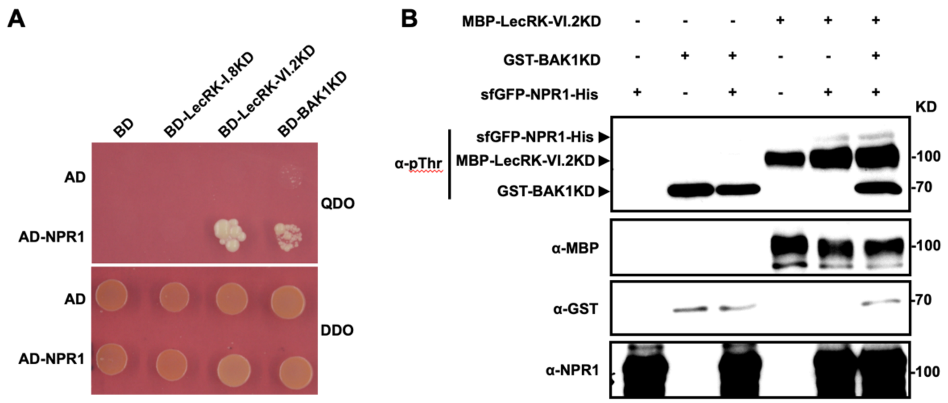

**Supplemental Figure 1 BAK1KD interacts with NPR1 but does not phosphorylate it *in vitro*.**

**(A)** Yeast two-hybrid assay examining the interaction between LecRK-1.8KD, LecRK-VI.2KD, or BAK1KD and NPR1. The fragments encoding the kinase domains (KDs) of LecRK-1.8, LecRK-VI.2, and BAK1 were cloned into the bait vector pGBKT7, and the full-length *NPR1* was cloned into the prey vector pGADT7. The bait and prey vectors were co-transformed into the yeast strain AH109, and the resulting yeast cells were grown on synthetic dextrose (SD) double dropout (DDO) (-Trp-Leu) medium. Interaction was determined by growth on SD quadruple dropout (QDO) (-Trp-Leu-His-Ade) medium. BD, DNA binding domain; AD, activation domain.

**(B)** Phosphorylation of NPR1 by LecRK-VI.2KD or BAK1KD *in vitro*. The kinase assay was performed by incubating MBP-LecRK-VI.2KD or GST-BAK1KD with sfGFP-NPR1-His as a substrate. Phosphorylation of NPR1 was detected using anti-pThr antibody. The abundance of the recombinant proteins was determined by immunoblotting with anti-MBP, anti-GST, and anti-NPR1 antibodies. The experiment was repeated twice with similar results.

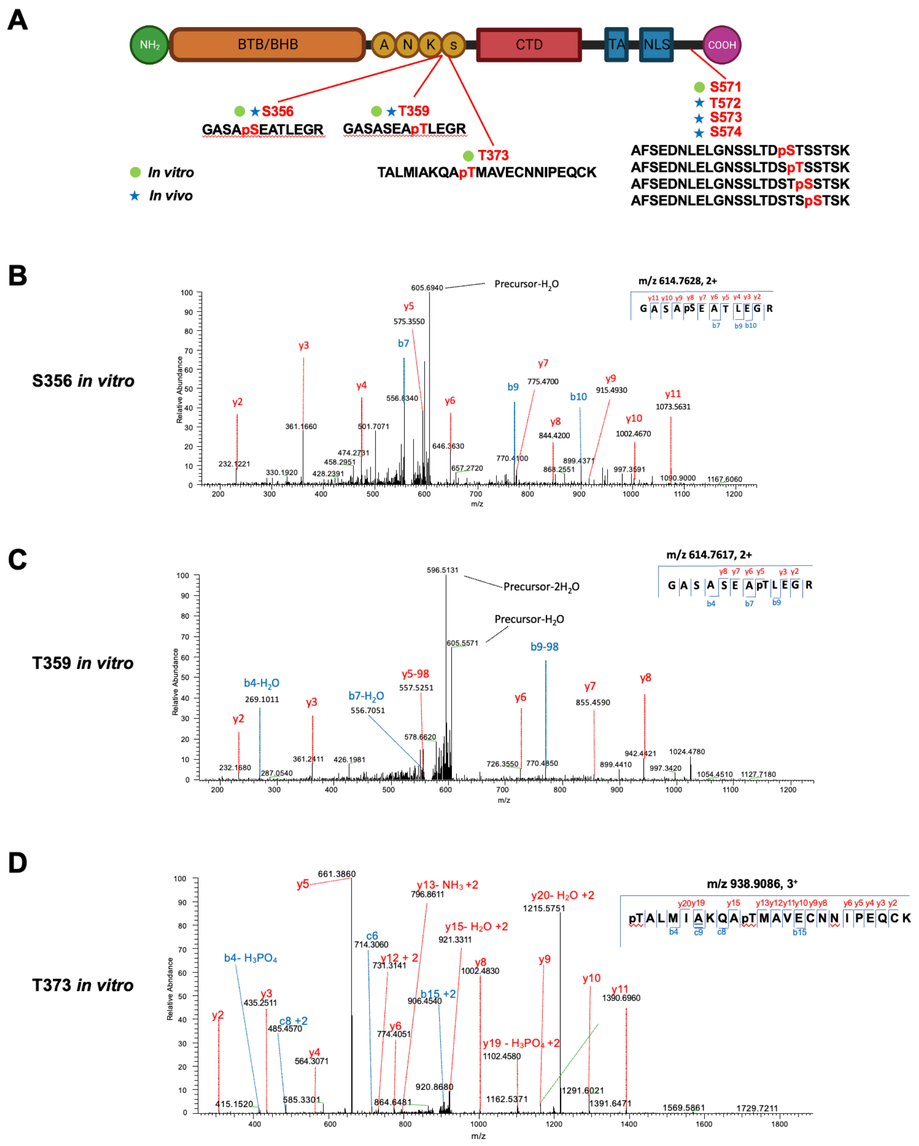

**Supplemental Figure 2 NPR1 *in vitro* phosphorylation sites identified by LC-MS/MS.**

**(A)** Schematic diagram of the NPR1 protein motifs with both *in vitro* and *in vivo* phosphorylation sites identified by LC-MS/MS.

**(B to D)** LC-MS/MS spectra of peptides harboring pS356 (**B**), pT359 (**C**), or pT373 (**D**) derived from sfGFP-NPR1-His incubated with MBP-LecRK-VI.2KD.

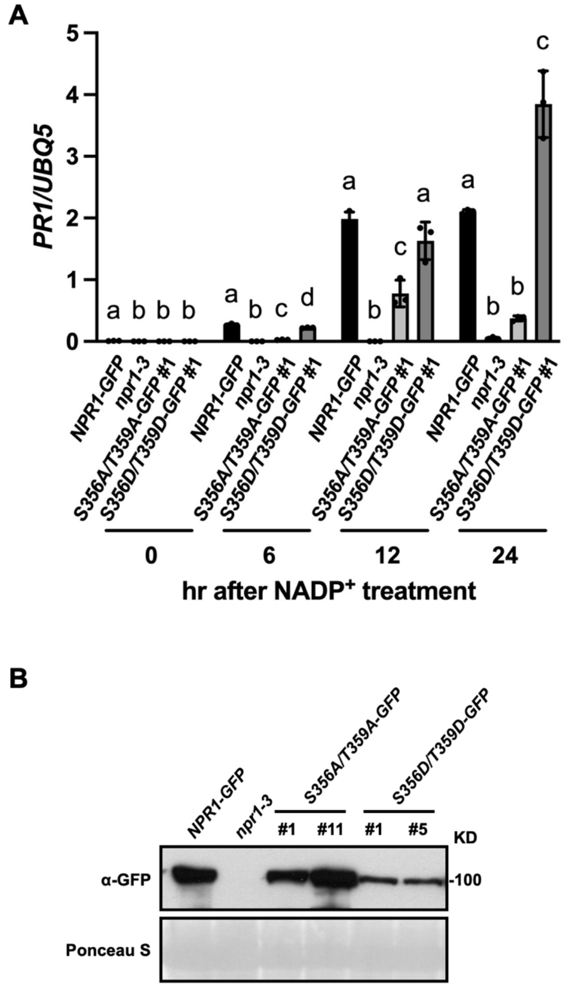

**Supplemental Figure 3 Phosphorylation of S356 and T359 is required for NADP$^+$-induced *PR1* gene expression.**

**(A)** NADP$^+$-induced *PR1* expression in the indicated genotypes. Leaves of four-week-old plants were infiltrated with 0.8 mM NADP$^+$. Leaf tissues were collected at the indicated time points for gene expression assay. Expression levels of *PR1* were normalized against the constitutively expressed *UBQ5*. Bars represent means ± SD (n = 3). Different letters denote significant differences ($p < 0.05$; one-way ANOVA). The comparison was made separately for each time point.

**(B)** Expression levels of the transgenic proteins in the indicated genotypes. Total proteins were analyzed by immunoblotting with anti-GFP antibody. Ponceau S staining of Rubisco was used as the loading control.

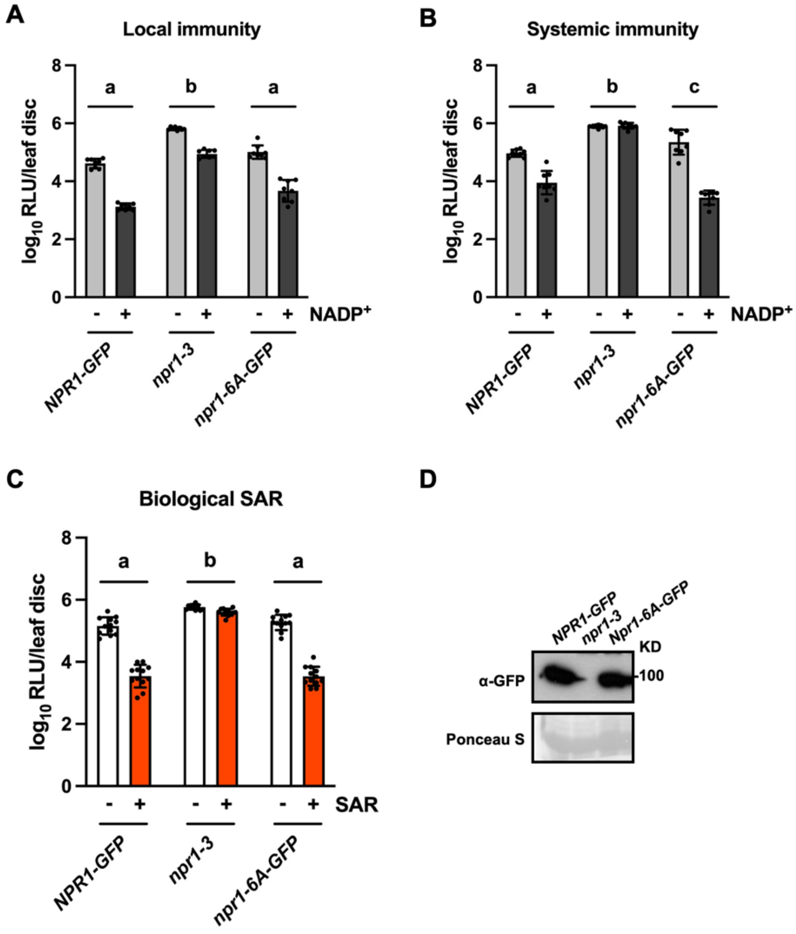

**Supplemental Figure 4 Phosphorylation of S/T571-576 is not required for NPR1's function in eNADP signaling and SAR.**

**(A and B)** $NADP^+$-induced local (**A**) and systemic (**B**) immunity in *npr1-3* and *npr1-3* complementation lines expressing *NPR1-GFP* or *npr1-6A-GFP*. *6A*, *S571A/T572A/S573A/S574A/T575A/S576A*. Two (**A**) or three leaves (**B**) on each plant were infiltrated with 0.8 m $NADP^+$ or water. Four hr later, either the infiltrated leaves (**A**) or two upper systemic leaves (**B**) were inoculated with *Psm-lux* ($OD_{600}$ = 0.001). Samples were taken 2.5 days later. Bars represent means ± SD (n = 8). Different letters denote significant differences ($p < 0.05$; two-way ANOVA).

**(C)** Biological induction of SAR in the indicated genotypes. Three lower leaves on each four-week-old plant were infiltrated with *Psm* ($OD_{600}$= 0.002) or 5 mM $MgCl_2$. Two days later, two systemic leaves were inoculated with a *Psm-lux* suspension ($OD_{600}$ = 0.001). Samples were collected three days later. Bars represent means ± SD (n = 12). Different letters denote significant differences ($p < 0.05$; two-way ANOVA).

**(D)** Expression levels of the transgenic proteins in the indicated genotypes. Total proteins were analyzed by immunoblotting with the anti-GFP antibody. Ponceau S staining of Rubisco was used as the loading control.

The experiments in (**A**-**C**) were repeated at least three times with similar results.

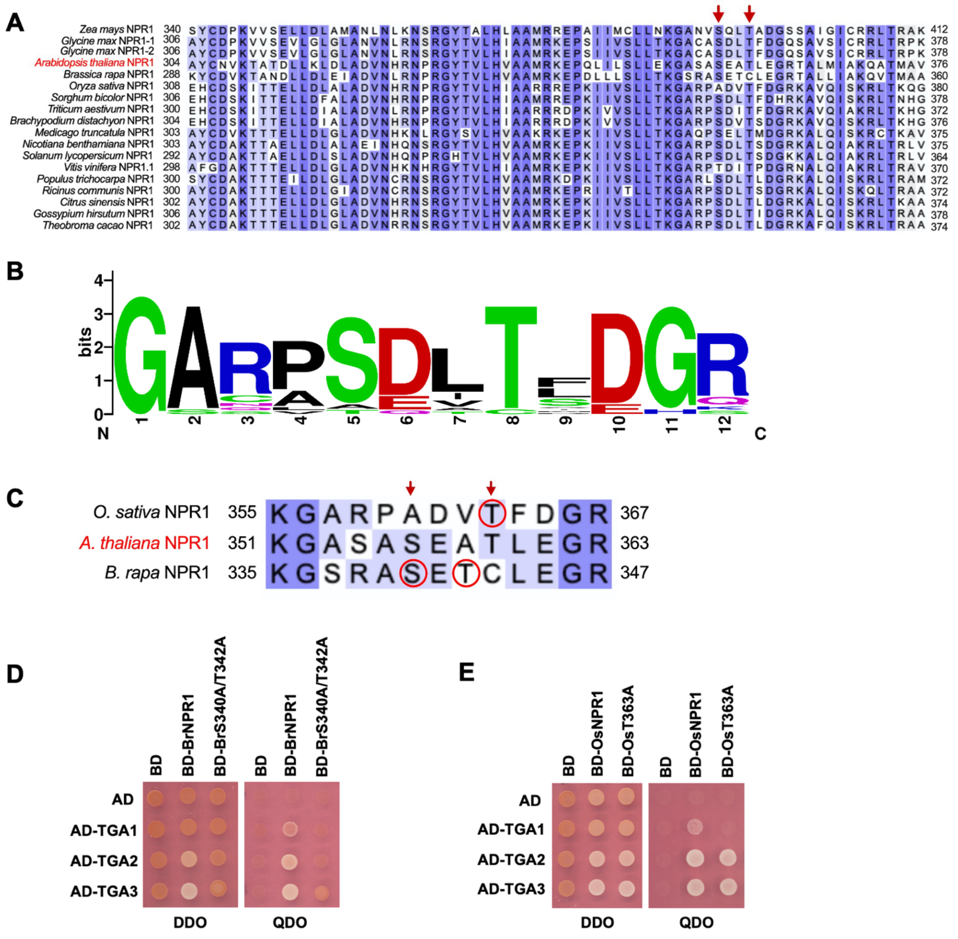

**Supplemental Figure 5 Conservation of the *Arabidopsis* NPR1 S356 and T359 phosphosites across plant species.**

**(A)** Multiple alignment of 18 NPR1 orthologs from 17 plant species using the “Align” tool in UniProt with default settings. The red arrows indicate the conserved Ser and Thr.

**(B)** WebLogo representation of the region surrounding the conserved Ser and Thr of 18 NPR1 orthologs from 17 plant species.

**(C)** The Ser/Thr residues (indicated by red arrows) corresponding to the *Arabidopsis* NPR1 S356 and T359 in *O. sativa* and *B. rapa* NPR1 orthologs. The Ser/Thr residues that were mutated in this study are marked with red circles.

(**D** and **E**) Yeast two-hybrid assay examining the interaction between TGA1, TGA2, or TGA3 and BrNPR1 (**D**), BrS340A/T342A (**D**), OsNPR1 (**E**) or OsT363A (**E**). The full-length *BrNPR1*, *BrS340A/T342A*, *OsNPR1*, and *OsT363A* were cloned into the bait vector pGBKT7, and the full-length *TGA1*, *TGA2*, and *TGA3* were in the prey vector pGADT7. The bait and prey vectors were co-transformed into the yeast strain AH109, and the resulting yeast cells were grown on DDO medium. Interaction was determined by growth on QDO medium. BD, DNA binding domain; AD, activation domain.

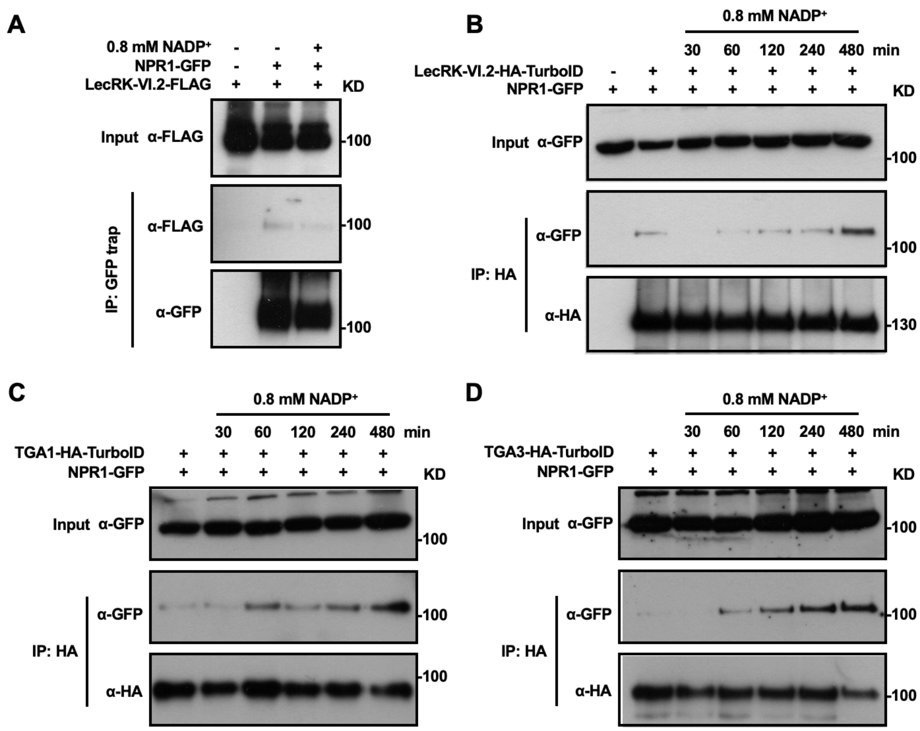

**Supplemental Figure 6 NADP$^+$ treatment triggers dissociation of NPR1 from LecRK-VI.2 and facilitates NPR1-TGA interaction.**

**(A)** NPR1-GFP and LecRK-VI.2-FLAG were transiently co-expressed in *N. benthamina*. Two days after agroinfiltration, the infiltrated leaves were treated with or without 0.8 mM NADP$^+$ for 30 min. Immunoprecipitation was performed using GFP-trap magnetic agarose beads. The precipitated proteins were analyzed by immunoblotting with anti-FLAG and anti-GFP antibodies.

**(B)** Co-immunoprecipitation assay of the interaction between NPR1 and LecRK-VI.2 *in vivo*. Leaves of four-week-old double transgenic plants expressing NPR1-GFP and LecRK-VI.2-HA-TurboID were treated with 0.8 mM NADP$^+$ for the indicated times. Immunoprecipitation was conducted using anti-HA magnetic beads. The precipitated proteins were analyzed by immunoblotting with anti-GFP and anti-HA antibodies. Transgenic plants expressing NPR1-GFP without $NADP^+$ treatment were included as negative controls.

**(C** and **D)** Co-immunoprecipitation assay of the interaction between NPR1-GFP and TGA1 (**C**) or TGA3 (**D**) *in vivo*. Leaves of four-week-old double transgenic plants expressing NPR1-GFP and TGA1-HA-TurboID or TGA3-HA-TurboID were treated with 0.8 mM $NADP^+$ for the indicated times. Immunoprecipitation was conducted using anti-HA magnetic beads. The precipitated proteins were analyzed by immunoblotting with anti-GFP and anti-HA antibodies. The molecular mass markers in kilo Daltons (KD) are indicated on the right, and the experiments were repeated at least twice with similar results.

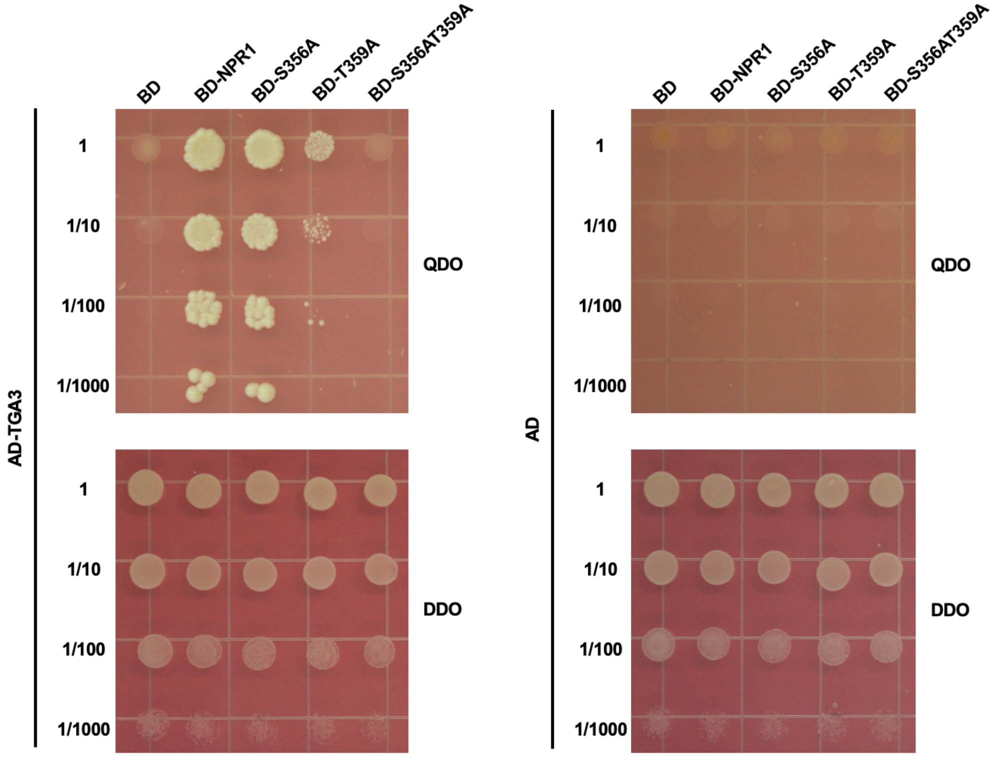

**Supplemental Figure 7 Phosphorylation of S356 and T359 additively contributes to NPR1-TGA3 interaction.**

Yeast two-hybrid assay examining the interaction between NPR1 or its phospho-mutants and TGA3. The full-length *NPR1* and its phospho-mutants were cloned into the bait vector pGBKT7, and the full-length *TGA3* was in the prey vector pGADT7. The bait and prey vectors were co-transformed into the yeast strain AH109, and the resulting yeast cells were grown on DDO medium. Interaction was determined by growth on QDO medium. BD, DNA binding domain; AD, activation domain.

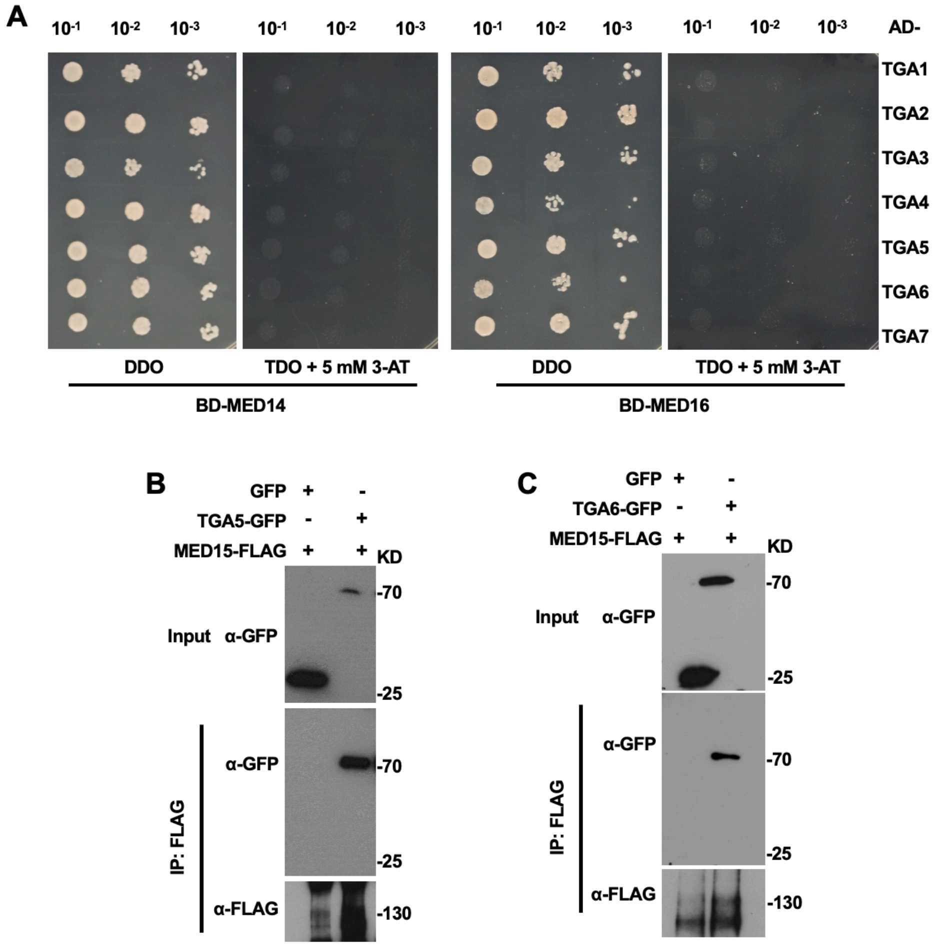

**Supplemental Figure 8 TGAs interact with MED15 but not MED14 and MED16**

**(A)** Yeast two-hybrid analysis of interaction between TGAs and MED14 or MED16. The full-length *MED14* and *MED16* were cloned into the bait vector pGBKT7, and the full-length *TGAs* were in the prey vector pGADT7. The bait and prey vectors were co-transformed into the yeast strain AH109, and the resulting yeast cells were grown on DDO medium. Interaction was determined by growth on TDO medium supplemented with 5 mM 3-AT. BD, DNA binding domain; AD, activation domain.

**(B** and **C)** Co-immunoprecipitation assays of the interactions between MED15 and either TGA5 (**B**) or TGA6 (**C**) in *N. benthamiana*. MED15-FLAG was transiently co-expressed with either TGA5-GFP, TGA6-GFP, or GFP in *N. benthamiana*. Immunoprecipitation was carried out using anti-FLAG antibody-coupled agarose beads. The precipitated proteins were analyzed by immunoblotting with anti-GFP and anti-FLAG antibodies.

The experiments in (**B** and **C**) were repeated twice with similar results.

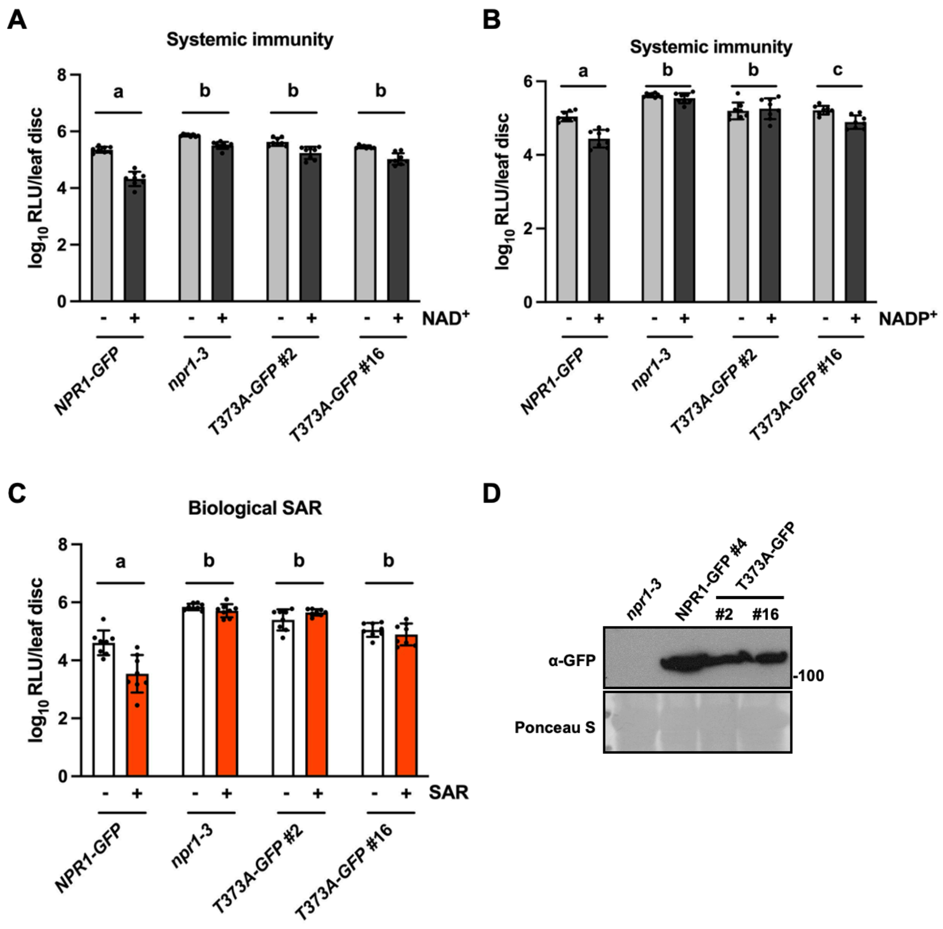

**Supplemental Figure 9 Phosphorylation of T373 is required for NPR1 function in eNADP signaling and SAR.**

**(A** and **B)** $NADP^+$-induced local (**A**) and systemic (**B**) immunity in *npr1-3* and *npr1-3* complementation lines expressing NPR1-GFP or T373A-GFP. Two (**A**) or three leaves (**B**) on each plant were infiltrated with 0.8 m $NADP^+$ or water. Four hr later, either the infiltrated leaves (**A**) or two upper systemic leaves (**B**) were inoculated with *Psm-lux* ($OD_{600}$ = 0.001). Samples were taken 2.5 days later. Bars represent means ± SD (n = 8). Different letters denote significant differences ($p < 0.05$; two-way ANOVA).

**(C)** Biological induction of SAR in the indicated genotypes. Three lower leaves on each four-week-old plant were infiltrated with *Psm* ($OD_{600}$= 0.002) or 5 mM $MgCl_2$. Two days later, two systemic leaves were inoculated with a *Psm-lux* suspension ($OD_{600}$ = 0.001). Samples were collected three days later. Bars represent means ± SD (n = 8). Different letters denote significant differences ($p < 0.05$; two-way ANOVA).

**(D)** Expression levels of the transgenic proteins in the indicated genotypes. Total proteins were analyzed by immunoblotting with the anti-GFP antibody. Ponceau S staining of Rubisco was used as the loading control.

The experiments in (**A-C**) were repeated three times with similar results.

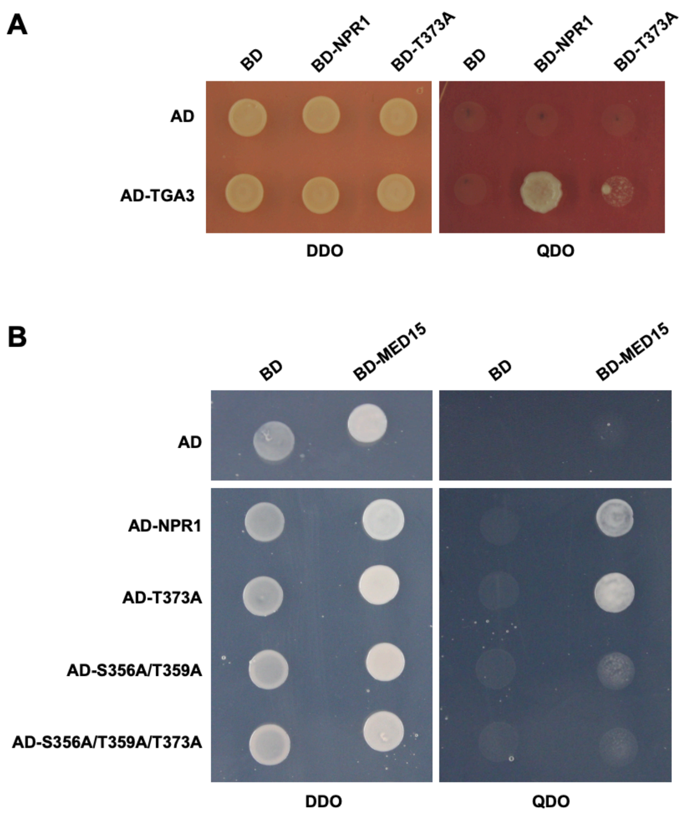

**Supplemental Figure 10 The effects of the T373A mutation on the NPR1 interactions with TGA3 and MED15.**

**(A)** Yeast two-hybrid assay examining the interaction between NPR1 or T373A and TGA3. The full-length *NPR1* and *T373A* were cloned into the bait vector pGBKT7, and the full-length *TGA3* was in the prey vector pGADT7.

**(B)** Yeast two-hybrid assay examining the interaction between MED15 and NPR1, T373A, S356A/T359A, or S356A/T359A/T373A. The full-length *MED15* was in pGBKT7, and the full-length *NPR1*, *T373A*, *S356A/T359A*, or *S356A/T359A/T373A* were cloned into pGADT7. The bait and prey vectors were co-transformed into the yeast strain AH109, and the resulting yeast cells were grown on DDO medium. Interaction was determined by growth on QDO medium. BD, DNA binding domain; AD, activation domain.

**Supplemental Table 1 Primers used in this study.**

| **Name** | **Sequence (5'-3')** | **Method** |
|---|---|---|
| **Cloning** | | |
| **pCAMBIA-1300S-GFP** | | |
| *NPR1-SacI-F* | TCTCGAGCTTTCGCGAGCTCATGGACACCACCATTGATGGATTC | |
| *NPR1-SalI-R* | CTTTACCCATGTCGACCCGACGACGATGAGAGAGTTTACG | in-fusion |
| *TGA1-BamHI-F* | CGCGGATCCATGAATTCGACATCGACACATTTT | |
| *TGA1-SalI-R* | ACGCGTCGACCGTTGGTTCACGATGTCGAGT | ligation |
| *TGA2-BamHI-F* | CGCGGATCCATGGCTGATACCAGTCCGAGAA | |
| *TGA2-SalI-R* | ACGCGTCGACCTCTCTGGGTCGAGCAAGCC | ligation |
| *TGA3-BamHI-F* | CGCGGATCCATGGAGATGATGAGCTCTTCTTCTT | |
| *TGA3-SalI-R* | ACGCGTCGACAGTGTGTTCTCGTGGACGAGCT | ligation |
| *TGA5-BamHI-F* | CGCGGATCCATGGGAGATACTAGTCCAAGAACATCAG | |
| *TGA5-SalI-R* | ACGCGTCGACCTCTCTTGGTCTGGCAAGCCAT | ligation |
| *TGA6-BamHI-F* | CGCGGATCCATGGCTGATACCAGTTCAAGGACT | |
| *TGA6-SalI-R* | ACGCGTCGACCTCTCTTGGCCGGGCAAG | ligation |
| *OsNPR1-KpnI-F* | GCTTTCGCGAGCTCGGTACCATGGAACCACCCACATCACATGTG | |
| *OsNPR1-BamHI-R* | CACCTCTAGAGGATCCCCTTCTTGGTCTAATGGCTCC | in-fusion |
| *BrNPR1-KpnI-F* | GCTTTCGCGAGCTCGGTACCATGGATTCCTTTGCTGGATTCGG | |
| *BrNPR1-BamHI-R* | CACCTCTAGAGGATCCGCGACGCCGATGGTAGAG | in-fusion |
| **pCAMBIA-1300S-FLAG** | | |
| *NPR1-SacI-F* | TCTCGAGCTTTCGCGAGCTCATGGACACCACCATTGATGGATTC | |
| *NPR1-SalI-R* | CTTTACCCATGTCGACCCGACGACGATGAGAGAGTTTACG | In-fusion |
| *MED15-KpnI-F* | CGGGGTACCATGGATAATAACAATTGGAGGCCTT | |
| *MED15-SalI-R* | GCGTCGACGGAAGCTGCTACATATTTCTCCCAA | ligation |
| **pGBKT7** | | |
| *LecRK-VI.2KD-NdeI-F* | GGAATTCCATATGTTCTTCGTCATGTACAAGAAGAG | |
| *LecRK-VI.2KD-XhoI-R* | CCCTCGAGCTACTGACTGATACGAGAAGTC | ligation |
| *LecRK-I.8KD-NDeI-F* | GGAATTCCATATGCTATACAGAAGAAACAAGTATGCAG | |
| *LecRK-I.8KD-SalI-R* | GCGTCGACTCATCGTCCAATTCCGTATTG | ligation |
| *BAK1KD-EcoRI-F* | GGAATTCCGAAGGAAAAAGCCGCAGGAC | |
| *BAK1KD-SalI-R* | GCGTCGACTTATCTTGGACCCGAGGGGTA | ligation |
| *OsNPR1-NdeI-F* | AGGAGGACCTGCATATGATGGAACCACCCACATCACATGTG | |
| *OsNPR1-BamHI-R* | GCAGGTCGACGGATCCTCACCTTCTTGGTCTAATGGCTCC | in-fusion |
| *BrNPR1-NdeI-F* | AGGAGGACCTGCATATGATGGATTCCTTTGCTGGATTCGG | |
| *BrNPR1-BamHI-R* | GCAGGTCGACGGATCCTCAGCGACGCCGATGGTAGAG | in-fusion |
| *MED14-BamHI-F* | CGGGATCCATGGCGGAATTAGGGCAACAG | |
| *MED14-XhoI-R* | CCCTCGAGCTATATGGTAAATTCCTTTTGATG | ligation |
| *MED15-NdeI-F* | GGAATTCCATATGGATAATAACAATTGGAGGCCTT | |
| *MED15-BamHI-R* | CGGGATCCTCAGGAAGCTGCTACATATTTCTCC | ligation |
| *MED16-EcoRI-F* | GGAATTCATGAATCAGCAAAACCCAGAAGA | |
| *MED16-SalI-R* | GCGTCGACCTATACAACACGGACCCACGTTCC | ligation |
| **pGADT7** | | |
| *NPR1-NdeI-F* | CAGATTACGCTCATATGATGGACACCACCATTGATGGATTCG | |
| *NPR1-BamHI-R* | CGAGCTCGATGGATCCTCACCGACGACGATGAGAGAG | in-fusion |
| **pCAMBIA1300S-NlucC-FLAG** | | |
| *NPR1-SacI-F* | CTCTCGAGCTTTCGCGAGCTCATGGACACCACCATTGATGGAT | |
| *NPR1-BamHI-R* | TCCGCCGCTTCCGCCGGATCCCCGACGACGATGAGAGAGTTTA | in-fusion |
| *GUS-BamHI-F* | CGGTACCCGGGGATCCATGTTACGTCCTGTAGAAACCCC | |
| *GUS-BamHI-R* | CGCTTCCGCCGGATCCTTGTTTGCCTCCCTGCTGC | In-fusion |
| **pBASTA-HA-TurboID** | | |
| *Lti6b-XhoI-F* | CCCTCGAGATGAGTACAGCCACTTTCGTAG | |
| *Lti6b-XbaI-R* | GCTCTAGACTTGGTGATGATATAAAGAGCG | ligation |
| *LecRK-VI.2-XhoI-F* | CCCTCGAGATGGGCACACAAAGATCCATG | |
| *LecRK-VI.2-XbaI-R* | GCTCTAGACTGACTGATACGAGAAGTCGAAGAAAC | ligation |
| *TGA1-KpnI-F* | CGGGGTACCATGAATTCGACATCGACACATTTT | |
| *TGA1-SalI-R* | GCGTCGACCGTTGGTTCACGATGTCGAGT | ligation |
| *TGA2-KpnI-F* | CGGGGTACCATGGCTGATACCAGTCCGAGAA | |

| | | |
|---|---|---|
| *TGA2-SalI-R* | GCGTCGACCTCTCTGGGTCGAGCAAGC | ligation |
| *TGA3-KpnI-F* | CGGGGTACCATGGAGATGATGAGCTCTTCTTCTT | |
| *TGA3-SalI-R* | GCGTCGACAGTGTGTTCTCGTGGACGAGCT | ligation |
| *LBD16-Kpn1-F* | GGGGTACCATGGCATCTTCCGGTAACGGT | |
| *LBD16-BamH1-R* | CGGGATCCGTTCTTCATCATTCTAAGAGCCAAAGCC | ligation |
| *mCherry-Kpn1-F* | CGGGGTACCATGGTTTCAAAAGGTGAAGAAGATAA | |
| *mCherry-BamH1-R* | CGCGGATCCTTTGTAGAGTTCATCCATTCCACC | ligation |
| *MED15-KpnI-F* | CGGGGTACCATGGATAATAACAATTGGAGGCCTT | |
| *MED15-SalI-R* | GCGTCGACGGAAGCTGCTACATATTTCTCCCAA | ligation |
| **pET28a-sfGFP-His** | | |
| *NPR1-BamHI/BsaI-F* | CGGGATCCGAGACCGACACCACCATTGATGGATTC | |
| *NPR1-SalI/BsaI-R* | ACGCGTCGACGAGACCCCGACGACGATGAGAGAGTTTAC | ligation |
| *sfGFP-NcoI-F* | CATGCCATGGTTAGCAAAGGTGAAGAAC | |
| *sfGFP-BamHI/EcoRI/SacI/SalI-R* | GCGTCGACGAGCTCGAATTCGGATCCGCTGCCTTTATACAGTTCATC | ligation |
| **Site-directed mutagenesis** | | |
| *npr1-S356A-F* | GAAAAAGGTGCAAGTGCAGCAGAAGCAACTTTGGAAG | |
| *npr1-S356A-R* | CTTCCAAAGTTGCTTCTGCTGCACTTGCACCTTTTTC | |
| *npr1-T359A-F* | GCAAGTGCATCAGAAGCAGCTTTGGAAGGTAGAACCGC | |
| *npr1-T359A-R* | GCGGTTCTACCTTCCAAAGCTGCTTCTGATGCACTTGC | |
| *npr1-S356A/T359A-F* | GAAAAAGGTGCAAGTGCAGCAGAAGCAGCTTTGGAAGGTAGAACCGC | |
| *npr1-S356A/T359A-R* | GCGGTTCTACCTTCCAAAGCTGCTTCTGCTGCACTTGCACCTTTTTC | |
| *npr1-S356D/T359D-F* | GTGCAAGTGCAGACGAAGCAGACTTGGAAGGTAGAACCGCACT | |
| *npr1-S356D/T359D-R* | AGTGCGGTTCTACCTTCCAAGTCTGCTTCGTCTGCACTTGCAC | |
| *npr1-S571A/T572A/S573A/S574A/T575A/S576A-F* | GGAAATTCGTCCCTGACAGATGCGGCTGCTGCCGCAGCGAAATCAACCGGTGGAAAG | |
| *npr1-S571A/T572A/S573A/S574A/T575A/S576A-R* | CTTTCCACCGGTTGATTTCGCTGCGGCAGCAGCCGCATCTGTCAGGGACGAATTTCC | |
| *LecRK-VI.2-K395E-F* | ATTCGGATCCCATCGCAGTGGAGAAGATAATTCCAAGTAGCAGGCAAG | |
| *LecRK-VI.2-K395E-R* | CTTGCCTGCTACTTGGAATTATCTTCTCCACTGCGATGGGATCCGAAT | |
| *Osnpr1-T363A-F* | AGCACGACCCGCGGATGTGGCTTTTGATGGAAGAAAAGCAGTGCA | |
| *Osnpr1-T363A-R* | TGCACTGCTTTTCTTCCATCAAAAGCCACATCCGCGGGTCGTGCT | |
| *Brnpr1-S340A/T342A-F* | GACAAAAGGGTCACGTGCAGCGGAAGCGTGTTTGGAAGGTAGAAC | |
| *Brnpr1-S340A/T342A-R* | GTTCTACCTTCCAAACACGCTTCCGCTGCACGTGACCCTTTTGTC | |
| **qPCR** | | |
| *PR1-F* | CTCATACACTCTGGTGGG | |
| *PR1-R* | ATTGCACGTGTTCGCAGC | |
| *UBQ5-F* | TCTCCGTGGTGGTGCTAAG | |
| *UBQ5-R* | GAACCTTTCCAGATCCATCG | |
